# The Hitchdock Domain in Kinesin-2 Tail Enables Adaptor Assembly and Cargo Binding

**DOI:** 10.1101/2025.03.21.644525

**Authors:** Xuguang Jiang, Radostin Danev, Baichun Niu, Sumio Ohtsuki, Haruaki Yanagisawa, Nobutaka Hirokawa, Masahide Kikkawa

## Abstract

Intracellular transport relies on motor proteins like kinesins to deliver essential cargoes along microtubules, yet the mechanisms of cargo recognition remain unclear. Here, we present high-resolution cryo-electron microscopy structures of the heterotrimeric kinesin-2 complex (KIF3A/KIF3B/KAP3) bound to the adenomatous polyposis coli (APC) cargo. Our findings reveal a previously uncharacterized KIF3 tail motif, termed the “Hitchdock domain,” which plays a pivotal role in mediating interactions with both the KAP3 adaptor and the APC cargo. In this domain, the KIF3A helical regions facilitate specific cargo binding, while the β-hairpin region and KIF3B provide structural support. Mutagenesis and molecular dynamics simulations confirm the domain’s functional importance. Interestingly, the Hitchdock/KAP3 structure suggests a conserved structural basis for cargo recognition across molecular motors, including kinesin-1 and dynein, which utilize similar hook-like architectures, highlighting the potential universality of this mechanism. Furthermore, our findings provide insights into kinesin-2 cargo specificity and offer a molecular framework for understanding related diseases.

## Introduction

The molecular mechanisms orchestrating the dynamic regulation of motor protein-driven intracellular transport have been a focal point of scientific inquiry for several decades ^1–4^. A key class of motor proteins, the kinesin superfamily (KIFs), plays a pivotal role in microtubule-based intracellular trafficking by recognizing and transporting various cargoes essential for diverse cellular processes ^5^. These kinesins are not only responsible for the movement of cargoes but also orchestrate complex regulatory mechanisms that ensure precise cargo delivery and cellular function ^4^.

Despite substantial progress in understanding kinesin’s interaction with microtubules, the molecular basis for kinesin-mediated cargo recognition and binding remains an unresolved question ^4,6^. Kinesins typically interact with cargoes through adaptors or accessory proteins that mediate specificity and control ^5,7^. However, due to the disordered nature of the kinesin tail region, which imparts significant structural flexibility, understanding the molecular mechanisms underlying kinesin-mediated transport, especially the interaction between kinesins, their adaptors, and cargoes, has been challenging.

Recent SAXS and negative-stain EM studies have revealed low-resolution binding models of kinesins and adaptors ^8,9^, and crystal studies of kinesin-1 light chain and cargo peptides have provided valuable insights into direct kinesin-cargo interactions ^10–12^, but these insights remain limited.

Kinesin-2 (KIF3/KAP3 in mammals) is one of the most widely expressed and abundant kinesins ^13–15^, forming a complex with heterodimeric KIF3 motors (KIF3A/KIF3B or KIF3A/KIF3C) and the armadillo repeat-containing adaptor KAP3 ^16–18^. Beyond its essential role in intraflagellar transport ^15,19,20^, KIF3/KAP3 has also been implicated in transporting diverse intracellular cargos, such as vesicles containing N-cadherin and NR2A subunit of NMDA receptor ^21–28^. Notably, the tumor suppressor protein adenomatous polyposis coli (APC) has been identified as a KIF3/KAP3 cargo involved in RNA transport crucial for neuronal development ^29,30^, accounting for ∼12% of KIF3-associated organelles ^31^. However, the mechanism of KIF3/KAP3 cargo recognition remains largely unclear.

To bridge this gap, we employed cryo-electron microscopy (cryo-EM) to obtain high-resolution structural analysis of the kinesin-2-adaptor-cargo (KIF3A/B-KAP3-APC) complex ^9,15–18,29–31^, providing the first detailed structural insight into the kinesin-adaptor-cargo system. This structural breakthrough reveals how the kinesin tail regions of KIF3A and KAP3 adaptor cooperate to mediate cargo recognition, while also highlighting isoform-specific differences in kinesin function, particularly within kinesin-2 families.

More importantly, our findings uncover a unique structural feature, the Hitchdock domain with Helix-β-hairpin-Helix motifs, in the kinesin-2 tail that is crucial for interaction with both the KAP3 adaptor and cargo APC. Supported by mutation experiments and molecular dynamics simulations, these results provide strong evidence for the functional relevance of this region. In the binding of both KAP3 and APC, the helices of the Hitchdock domain in KIF3A play a key role in the specific interaction, while the β-hairpin region and KIF3B provide structural support. Moreover, the discovery of the hook-like kinesin-2 Hitchdock-KAP3 structure suggests a conserved mechanism among molecular motors, including kinesin-1 and dynein, which utilize similar hook-like structures for cargo recognition. Notably, the structural similarities between the dynein FTS–HOOK–FHIP1B complex ^32,33^ and the kinesin-1 heavy chain/light chain cargo-binding region ^8^ with the KIF3 tail/KAP3 structure highlight the potential universality of this mechanism, warranting further investigation. The discovery of the conserved sequence motif for direct cargo recognition in the KIF3 Hitchdock region that directly interacts with the cargo facilitates further exploration of specific cargos and understanding of the pathogenesis of kinesin-2-related diseases.

## Results

### Structural Organization and Interfaces of the KIF3 C-terminus/KAP3 Complex

To understand how the tail region of the KIF3A/B heterodimer interacts with the cargo adaptor KAP3, we determined the high-resolution structure of the KIF3/KAP3 cargo-binding domain. Due to the elongated shape of the complex, we observed severe orientation bias, which was overcome by collecting an extensive dataset to enrich protein particles in diverse orientations. Data processing yielded two cryo-EM maps (2.83 and 2.75 Å resolution) representing distinct conformations (Supplementary Figure 1 and Supplementary Table 1). The maps shared highly similar regions, with Class 1 containing an additional density. Thus, we selected Class 1 for de novo model building using ModelAngelo ^34^, followed by manual refinement (Supplementary Figure 2). The final model resolved the core region of the ternary complex, encompassing KIF3A (residues 574–658), KIF3B (residues 580–675), and the Armadillo repeat (ARM) domain of KAP3 (residues 129–689) (Figure 1a-c).

**Figure 1.**
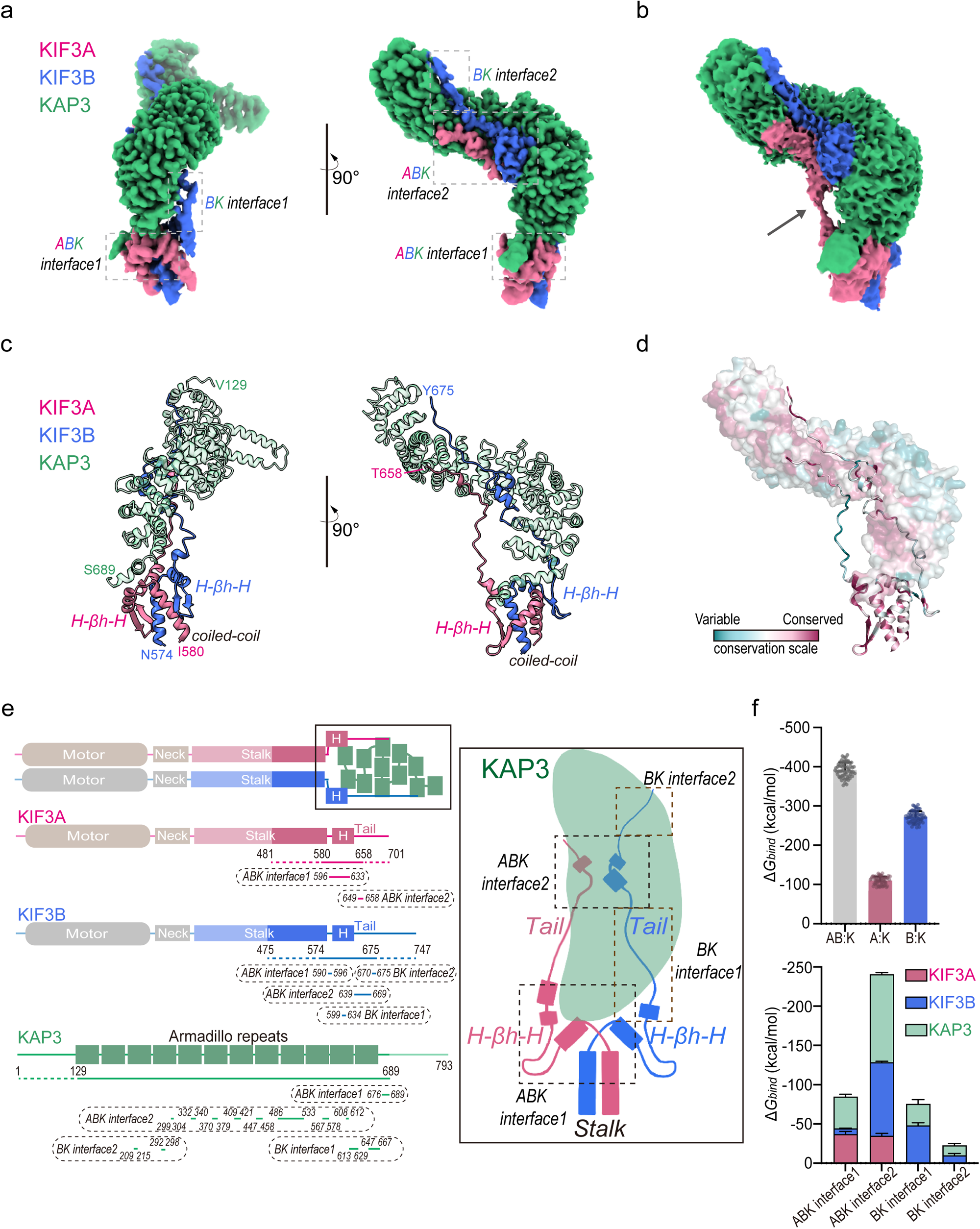
Basic architecture of the KIF3 C-terminus/KAP3 complex. (a) High-resolution cryo-EM map of the KIF3 C-terminal/KAP3 complex. Densities for KIF3A, KIF3B, and KAP3 are colored in red, blue, and green, respectively. The interface regions among the four molecules are indicated by dashed-line boxes, including two shared interfaces (ABK interfaces 1 and 2) and two KIF3B-specific interfaces (BK interfaces 1 and 2). (b) Volume of the KIF3A tail linking loop. A segment of the KIF3A linking loop density is visible in the raw EM map at a higher contour threshold. (c) Reconstructed structural model of the KIF3 C-terminus/KAP3 complex. The model includes partial stalk and tail regions of KIF3A/B (KIF3A residues 580–658; KIF3B residues 574–675) and the ARM domain of KAP3 (residues 129–689). The KIF3 heterodimer structure comprises the coiled-coil region at the stalk terminus, the helix-beta-hairpin-helix (H-βh-H) motif, and the extended tail region. (d) Conservation analysis of the KIF3 C-terminal/KAP3 structural model. Conservation scores were calculated using the ConSurf server (https://consurf.tau.ac.il/). (e) 2D domain organization of KIF3A/B/KAP3 and a cartoon diagram illustrating the binding mode between the KIF3 tail and KAP3. In the 2D schematic, the transparent regions represent areas not included in the KIF3/KAP3 construct. For the construct region, KIF3A (residues 481-701), KIF3B (residues 475-747), and KAP3 (residues 1-693), the visible portions in the structure are shown with solid lines, and the invisible portions are indicated with dashed lines. The regions involved in each interface are marked below the construct lines. (f) Binding free energy calculations based on MD-MMPBSA. The left panel shows the total binding free energy between the KIF3A/B tail and KAP3 (AB:K), as well as the individual contributions of KIF3A and KIF3B (A:K and B:K). The right panel summarizes the binding free energy contributions of each interface and the individual components of the complex. Bar graphs indicate mean ± SD.

The reconstructed structure revealed four interaction interfaces between KIF3 and KAP3, including two tripartite interfaces involving both KIF3A and KIF3B (ABK interfaces 1 and 2) and two additional interfaces specific to KIF3B-KAP3 interactions (BK interfaces 1 and 2) (Figure 1a). At the proximal tail region, adjacent to the stalk, KIF3A and KIF3B form a helix–beta-hairpin–helix (H-βh-H) motif, which interacts with the C-terminal region of KAP3, constituting ABK interface1 (Figure 1a-c). Further along the KIF3A/B tail, an extended loop reaches toward both sides of KAP3, converging at ABK interface2. Notably, the linking loop of KIF3A does not interact with KAP3 (Figure 1b, arrow), whereas KIF3B engages in direct contact at BK interface1 (Figure 1b-c). At ABK interface2, both KIF3A and KIF3B form short helices that interact with the middle region of KAP3. Subsequently, KIF3B extends further toward the N-terminal region of KAP3, forming BK interface2.

Conservation analysis revealed that the residues at these binding interfaces are relatively well conserved, particularly at ABK interface1 (Figure 1d). Due to the inherent flexibility of certain regions, some segments were unresolved in the cryo-EM density map, preventing the modeling of KIF3A beyond ABK interface2 (residues 658–701), KIF3B beyond BK interface2 (residues 675–747), and the N-terminal segment of KAP3 (residues 1–129) (Figure 1c and 1e).

Further molecular dynamics (MD) simulations coupled with MMPBSA-based binding free energy analysis revealed that KIF3A and KIF3B contribute ∼28% and ∼72%, respectively, to the overall interaction with KAP3. Notably, KIF3A exhibits a stronger contribution at ABK interface1, whereas its involvement in ABK interface2 is relatively weaker (Figure 1f).

### H-***β***h-H motif in the KIF3 Tail Serves as a Core Scaffold for KAP3 Interaction

Interestingly, unlike previous predictions suggesting that the kinesin-2 tail region is entirely disordered, our structural model reveals a well-defined H-βh-H motif at the N-terminal tail region immediately following the stalk (Figure 2a, Supplementary Figure 3a). This motif consists of a shared α-helix, β-hairpin, and 3_10_-helix (α-βh-3_10_) in both KIF3A and KIF3B. The C-terminal end of the KIF3A stalk engages with the α-helix and β-hairpin structures of KIF3B. In contrast, the C-terminal end of KIF3B similarly interacts with the corresponding structures of KIF3A, collectively reinforcing structural stability (Supplementary Figure 3a). Notably, in KIF3A, an additional short α-helix is observed following the 3_10_-helix, forming an αA-βh-3_10_-αB motif (Figure 2a).

**Figure 2.**
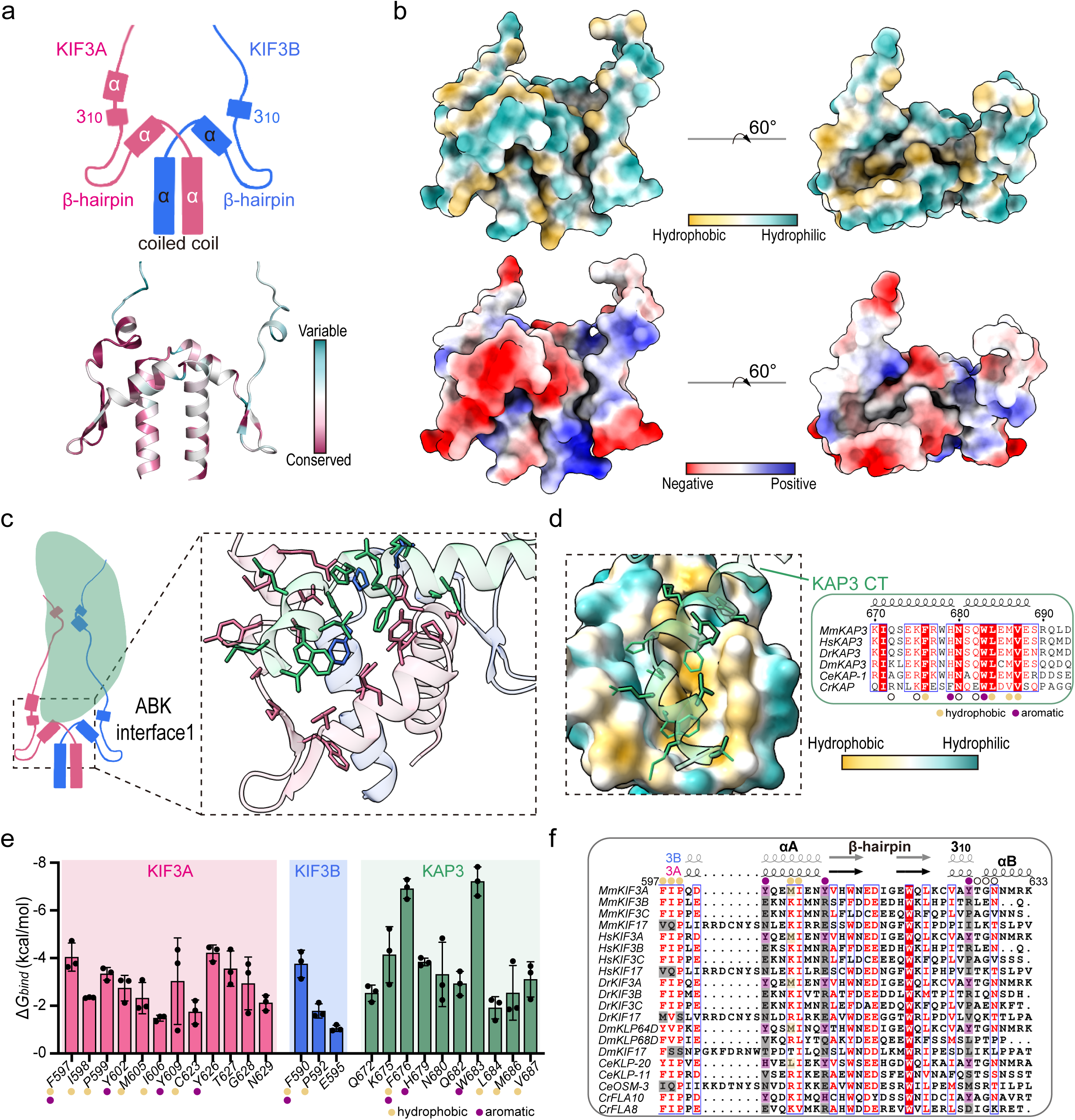
The stalk-tail junction of the KIF3 heterodimer forms a scaffold structure and binds KAP3. (a) Cartoon representation and conservation analysis of the stalk-tail junction of the KIF3 heterodimer. This region consists of a coiled-coil helix at the stalk terminus and an N-terminal helix-beta-hairpin-helix (H-βh-H) motif in the tail. Conservation analysis reveals that KIF3A is more conserved than KIF3B in this region (bottom panel). (b) Surface hydrophobicity and electrostatic properties of the scaffold region. (c) Binding mode of the scaffold region with KAP3 (ABK interface1). The C-terminal helix of KAP3 binds to the hydrophobic pocket on the KIF3A side, with minor contributions from KIF3B. (d) Insertion of the KAP3 C-terminal helix into the hydrophobic pocket of the KIF3 scaffold (left) and sequence alignment analysis (right). Key hydrophobic and aromatic residues in KAP3 (e.g., His679, Trp683, Leu684, Val697) are highly conserved across species. Yellow circles mark hydrophobic residues, purple circles mark aromatic residues, and open circles mark residues involved in other interactions. (e) Binding free energy contributions of residues involved in ABK interface1. Bar graphs indicate mean ± SD. (f) Structure-based sequence alignment of KIF3 residues involved in the binding interface. The alignment was generated using the ESPript server (https://espript.ibcp.fr/). Purple highlights indicate conserved residues in KIF3A critical for KAP3 binding, while gray highlights indicate non-conserved residues in KIF3B and KIF17.

Surface hydrophobicity and electrostatic analysis reveal that this motif in KIF3A forms a hydrophobic pocket on one side (Figure 2b, top-right panel), while its αA-βh surface is highly enriched with negative charges (Figure 2b, bottom-left panel). Within ABK interface1, the highly conserved C-terminal helix of KAP3 binds to this hydrophobic pocket (Figure 2c-d), primarily through hydrophobic and aromatic interactions with KIF3A (Figure 2d-e). Additionally, Lys675 in KAP3 further stabilizes the interaction via ionic interactions (Figure 2d). Moreover, Phe590 in KIF3B, located at the stalk-tail junction, also contributes to forming this hydrophobic pocket and enhances binding affinity. Notably, β-hairpin regions do not participate in KAP3 binding.

Sequence alignment analysis shows that the core residues forming the H-βh-H motif are highly conserved across the kinesin-2 family, including both heterodimeric (KIF3A/B/C) and homodimeric (KIF17) members (Figure 2f). This suggests that the motif is a shared structural feature within kinesin-2 tails. However, the residues directly involved in KAP3 binding are not universally conserved. Specifically, the hydrophobic and aromatic residues critical for KAP3 binding in KIF3A (Tyr602, Met605, Tyr609, and Tyr626) are conserved within KIF3A homologs but not in KIF3B/C (kinesin-2β) or KIF17 (kinesin-2γ, KAP3 indispensable) (Figure 2f). This suggests that KIF3A, the kinesin-2α subunit, plays a key role in KAP3 recognition. Furthermore, KIF17 lacks the conserved KIF3B/C residues (e.g., KIF3B Phe590) required for KAP3 interaction, further indicating that KIF17 does not bind KAP3.

In summary, the interaction between the KIF3 tail N-terminal region and the KAP3 C-terminal helix forms a stable structural foundation within the heterodimeric kinesin-2 tail. This interface may serve as a key determinant distinguishing heterodimeric and homodimeric kinesin-2 members in their ability to bind KAP3 and specific cargoes.

### KIF3 Tail Establishes Multiple Binding Sites with the Concave Surface of KAP3

Following the formation of the primary binding interface (ABK interface1) at the hydrophobic pocket of KIF3, the KIF3 tail extends further to establish a secondary binding interface, ABK interface2, with the concave surface of the KAP3 mid-region (Figure 3a). Unlike ABK interface1, where KAP3 C-terminal helix fits into a hydrophobic pocket of KIF3, at ABK interface2, both KIF3A and KIF3B adopt short helical conformations and interact with a highly conserved, electrostatically enriched, and partially hydrophobic concave surface of KAP3 (Figure 3a and Supplementary Figure 3b-c). KIF3A engages in ABK interface2 by forming a 3_10_-helix, anchoring to KAP3 primarily via hydrophobic interactions and hydrogen bonding. Notably, Tyr654, Leu655, Ala656, and Tyr657 contribute significantly to the binding, establishing up to five hydrogen bonds (Figure 3b-c). Meanwhile, the proximal segment of KIF3B tail adopts both a 3_10_-helix and an α-helix, where a cluster of seven positively charged residues facilitates electrostatic interactions with the negatively charged surface of KAP3 (Figure 3b). Among these, Arg654 forms a stable salt bridge with KAP3, playing a crucial role in stabilizing the complex (Figure 3a-c). Further, the distal segment of the KIF3B tail interacts with the hydrophobic patch on KAP3 (Figure 3a). Remarkably, KIF3B forms a total of 14 hydrogen bonds and two salt bridges with KAP3 (Supplementary Figure 3b-c), establishing an exceptionally stable binding interface. ABK interface2 engages a larger region and involves more extensive interactions (Figure 3 and Supplementary Figure 4), contributing approximately three times the binding energy compared to ABK interface1, as shown by MMPBSA free energy calculations (Figure 1f).

**Figure 3.**
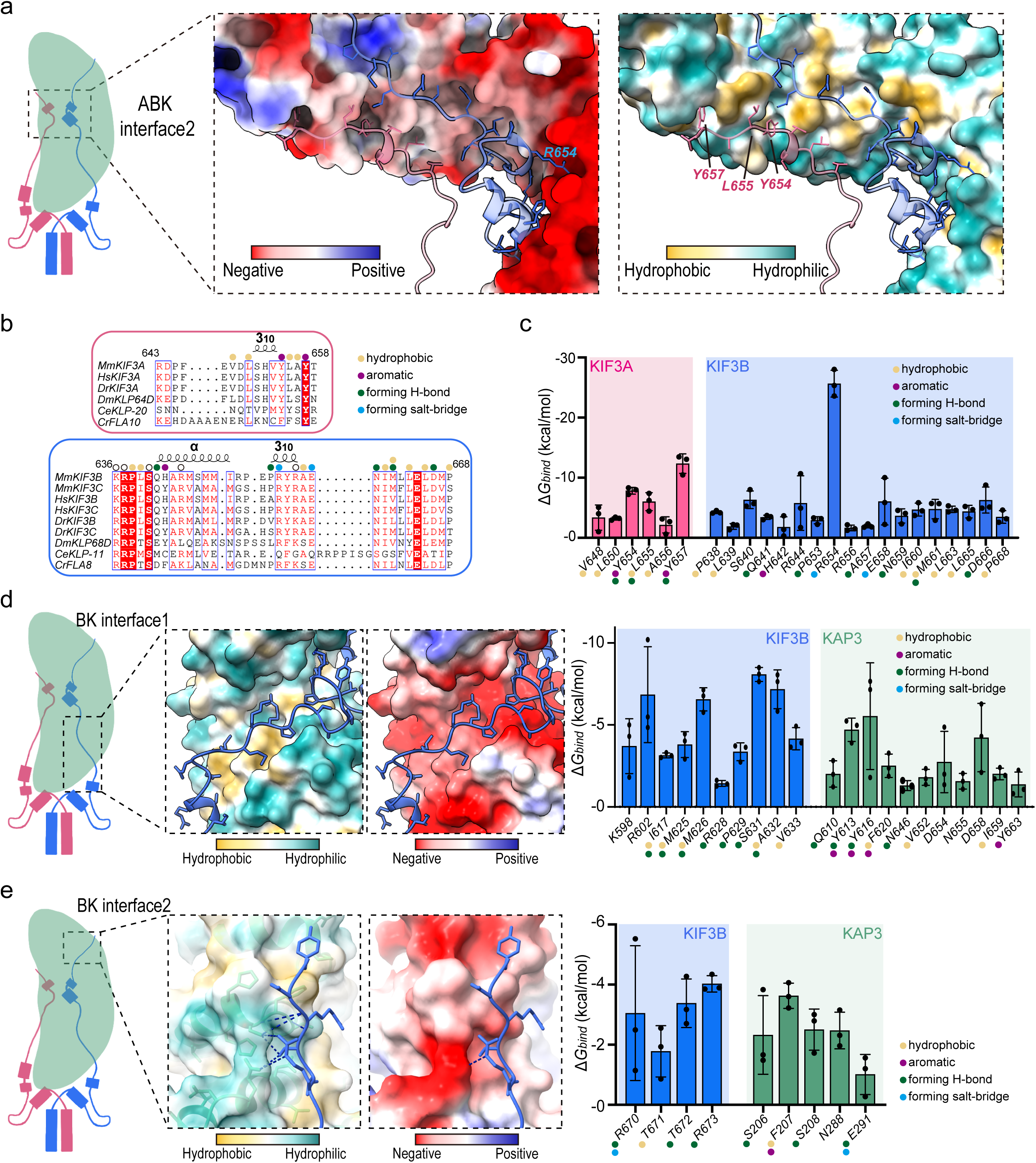
The KIF3 tail interacts with KAP3 at multiple sites. (a) The second binding interface (ABK interface2) on the concave surface of KAP3. The electrostatic (middle panel) and hydrophobic (right panel) properties of the KAP3 surface are shown, along with the cartoon structures of KIF3A/B segments involved in the interaction. Key interacting residues are labeled. (b) Sequence alignment of KIF3 residues involved in ABK interface2. Short helices formed by KIF3 and key interacting residues are highlighted. (c) Binding free energy contributions of residues involved in ABK interface2. Key interacting residues and interaction patterns are labeled. (d) The first KIF3B-specific binding interface (BK interface1). The electrostatic and hydrophobic properties of the KAP3 surface and the cartoon structure of the KIF3B segment are shown (left panel). Binding free energy contributions of residues involved in BK interface 1 are summarized (right panel). Key interacting residues are labeled. (e) The second KIF3B-specific binding interface (BK interface2). The electrostatic and hydrophobic properties of the KAP3 surface, the cartoon structure of the KIF3B segment, and the hydrogen bond network are shown (left panel). Binding free energy contributions of residues involved in BK interface2 are summarized (right panel). Key interacting residues are labeled. Bar graphs indicate mean ± SD.

Beyond ABK interfaces 1 and 2, KIF3B engages in two additional binding interfaces absent in KIF3A. BK interface1: A 10-residue segment following the H-βh-H motif in KIF3B interacts with the dorsal surface of KAP3 (Figure 3d); and BK interface2: A five-residue extension from the distal KIF3B tail interacts with the N-terminal region of KAP3 (Figure 3e). These additional binding sites contribute approximately one-fourth of the total binding energy (Figure 1f), further reinforcing the stability of the KIF3 tail/KAP3 complex.

Collectively, beyond the primary binding interface, the KIF3 tail interacts with KAP3 through multiple binding sites, engaging a highly conserved concave surface. This multi-site binding strategy provides a molecular basis for the stable association between KIF3 and its cargo adaptor, KAP3.

### APC Binds to the Core Scaffold Region of the KIF3 tail/KAP3 Complex

Having elucidated how the KIF3 tail stably associates with its adaptor protein KAP3, we next sought to understand how the KIF3 tail/KAP3 complex engages cargo proteins. To this end, we assembled a tetrameric KIF3 tail/KAP3-APC complex ^9,21,29,30,35^ using the validated KIF3/KAP3-binding ARM repeat region of APC and attempted single-particle cryo-EM reconstruction. However, the complex exhibited a severe orientation bias, making high-resolution map reconstruction challenging. By selectively reducing particles in the predominant orientation, we rebalanced the distribution and obtained an EM density map for the tetrameric complex (Supplementary Figure 5). Compared to the KIF3/KAP3 map, the KIF3/KAP3-APC map revealed an additional density at the ABK interface1 region, corresponding to the APC ARM domain (Figure 4a and Supplementary Figure 5).

**Figure 4.**
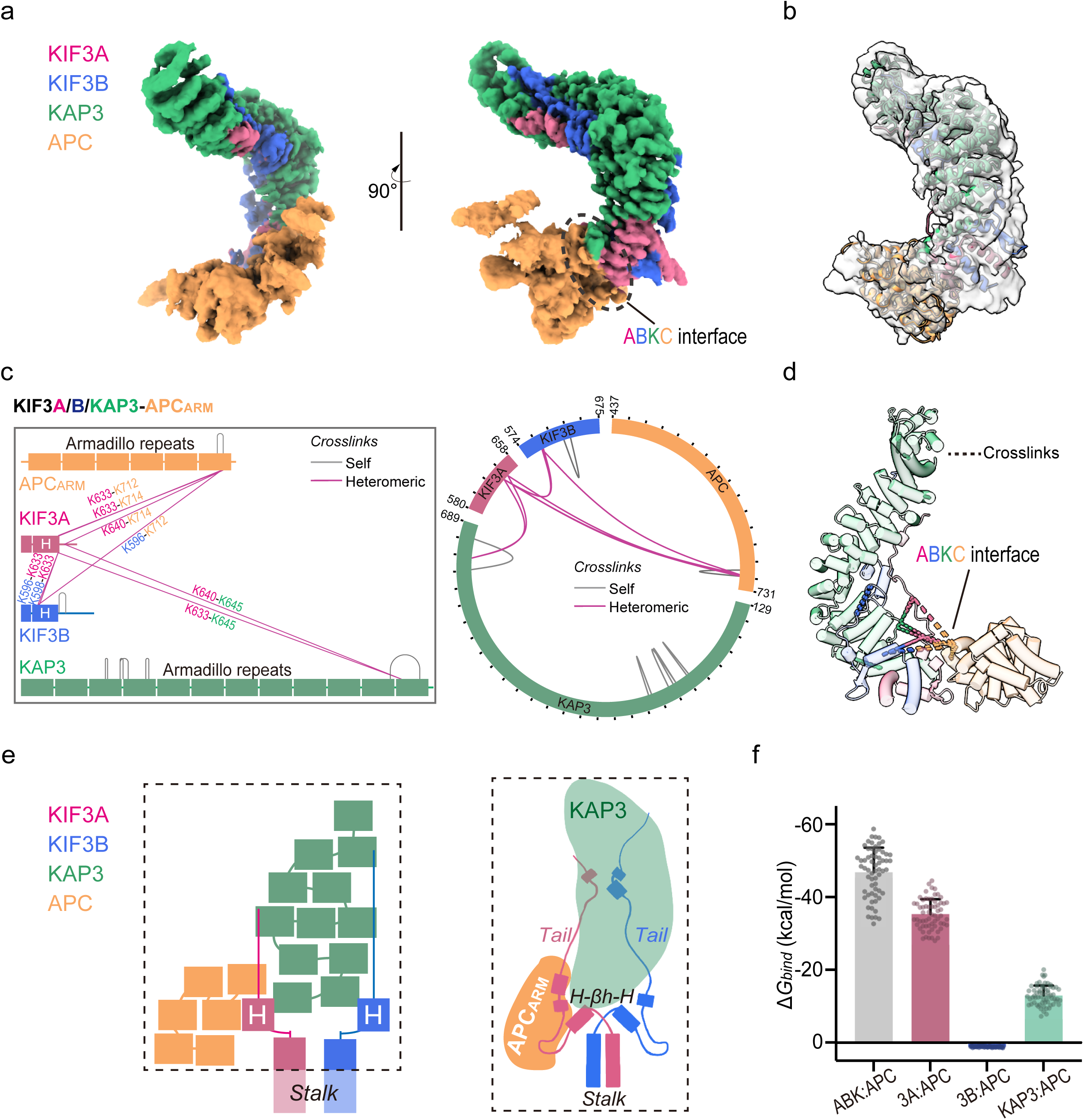
Structural determination and XL-MS analysis of the KIF3/KAP3-APC complex. (a) Cryo-EM map of the KIF3 C-terminal/KAP3-APC complex, with APC density colored in yellow. The binding interface between the KIF3/KAP3 and APC is labeled (ABKC interface). (b) Model of the KIF3 C-terminal/KAP3-APC complex. The cryo-EM map is shown in transparent gray, and the APC model is colored yellow. (c) 2D diagram (left) and circular representation (right) of XL-MS results for the KIF3 C-terminal/KAP3-APC complex. Intermolecular crosslinks are shown in purple, and intramolecular crosslinks are shown in gray. Only crosslinks visible in the structural model are displayed. Full data are available in Supplementary Figure 7 and Supplementary Table 2. (d) XL-MS results mapped onto the structural model. Intermolecular crosslinks are depicted using dashed lines in two colors, each corresponding to the respective components, while intramolecular crosslinks are represented by dashed lines in a single color matching the component. Crosslink pairs are within theoretical distances, and the ABKC interface is enriched with crosslinks, validating the structural model. (e) Domain organization of the KIF3A/B/KAP3-APC complex and a 2D diagram illustrating the binding mode. (f) Binding free energy calculations for KIF3/KAP3 binding to APC. The total binding free energy (ABK: APC) and individual contributions of KIF3A, KIF3B, and KAP3 (3A:APC, 3B:APC, and KAP3:APC) are shown. Bar graphs indicate mean ± SD.

While the KIF3/KAP3 region maintained an average resolution of ∼3 Å, the local resolution of the APC ARM domain was lower than 4 Å, making it unsuitable for de novo model building. To construct an initial model, we first docked the KIF3 tail/KAP3 structure, followed by integrating the available crystal structure of the APC ARM domain complex (PDB: 4YJE), using the raw map for docking to mitigate the effects of oversharpening. The model was further refined using MD-based flexible fitting and manual adjustments, improving its alignment with the EM density (Figure 4b, Supplementary Figure 6).

To validate the model, we conducted crosslinking mass spectrometry (XL-MS) analysis (Supplementary Figure 7 and Supplementary Table 2) and identified 15 lysine crosslinks within our model—seven intramolecular and eight intermolecular (Figure 4c and Supplementary Figure 7b). All detected lysine pairs were within the spatial constraints of the BS3 crosslinker, supporting the structural model, particularly the ABK-APC binding region (ABKC interface) (Figure 4d).

Based on the cryo-EM and XL-MS results, we found unexpectedly that the C-terminal ARM repeats of APC bind adjacent to the KIF3A tail and the C-terminal region of KAP3, specifically at the ABK interface1 region (Figure 4e). Further MD simulations (Supplementary Figure 8) revealed that KIF3A/APC interactions provided the dominant binding energy, three times that of KAP3/APC, while KIF3B did not contribute to APC binding (Figure 4f).

### KIF3A and KAP3 Cooperatively Bind APC via a Hydrophobic Pocket

To elucidate the molecular mechanism of APC ARM domain recognition by the KIF3/KAP3 complex, we performed a detailed structural analysis on the ABKC interface. The interaction site primarily involves the βh-3_10_-αB segment within the motif (αA-βh-3_10_-αB) of KIF3A, which contributes 11 residues, and the C-terminal helix of KAP3, which engages three residues, collectively interacting with 17 residues of the APC ARM domain (Figure 5a). These interactions are predominantly mediated by hydrophobic contacts and hydrogen bonding.

**Figure 5.**
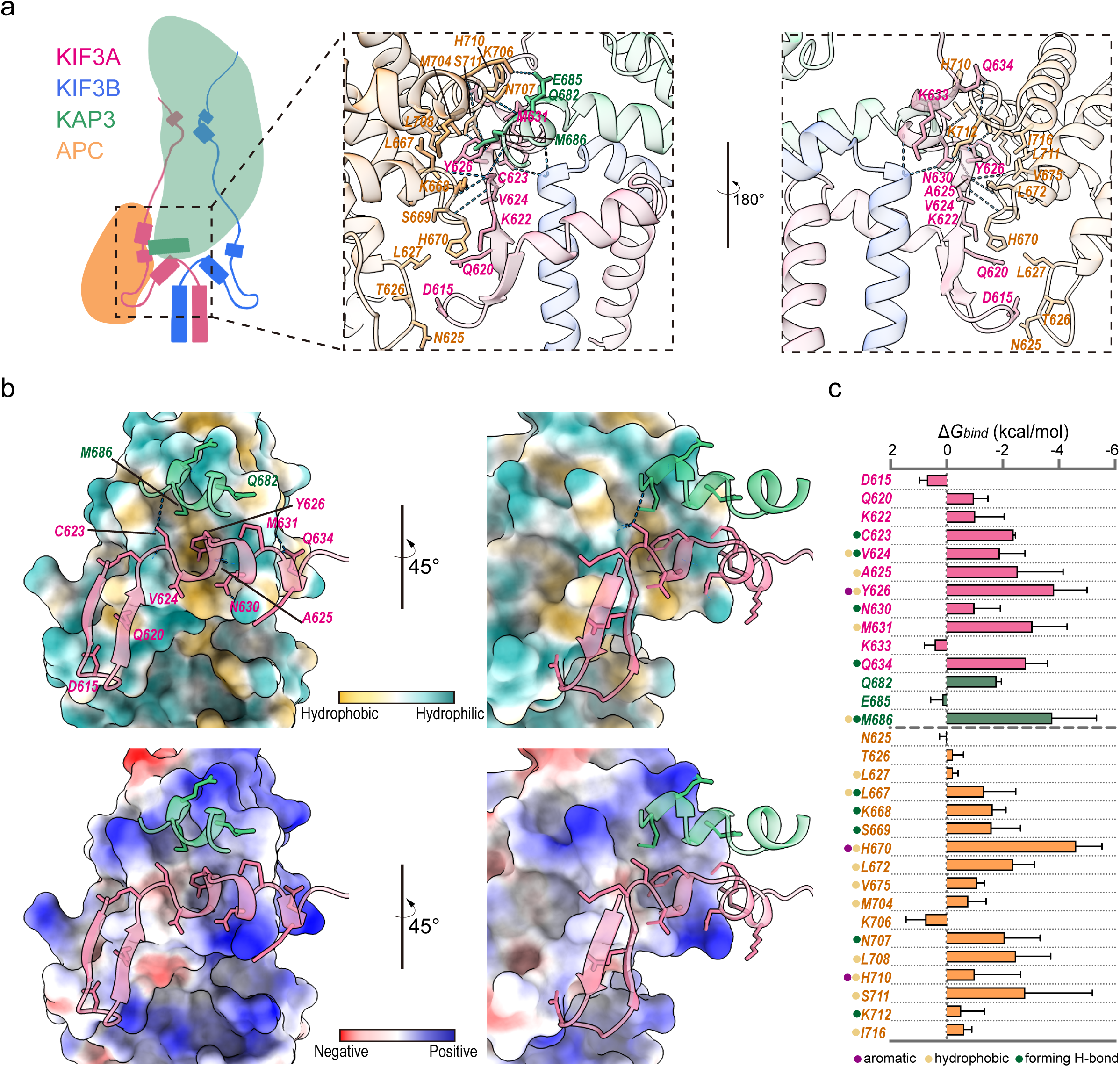
Cooperative binding of APC by KIF3A and KAP3. (a) Cartoon representation of the KIF3A/KAP3-APC binding interface (ABK interface1). The H-βh-H motif of KIF3A and the C-terminal helix of KAP3 jointly bind the ARM domain of APC. Key interacting residues and hydrogen bonds are labeled (right panel). (b) Hydrophobic (upper panel) and electrostatic (bottom panel) properties of the APC surface, along with the cartoon structures of KIF3A/KAP3 segments involved in the interaction. The interface is primarily hydrophobic, with key interacting residues labeled. (c) Binding free energy contributions of residues involved in the KIF3A/KAP3-APC binding interface. Key interacting residues and interaction patterns are labeled. Bar graphs indicate mean ± SD.

Within this interface, the 3_10_-αB region of KIF3A plays a central role, with Val624, Ala625, and Tyr626 from the 3_10_-helix inserting into a hydrophobic pocket formed at the C-terminal region of APC ARM. Additionally, Cys623 and Asn630 contribute to stabilization through four hydrogen bonds with APC (Figure 5a-b). Further interaction is mediated by Met631 and Gln684 from the αB helix, which establishes additional contacts with adjacent APC residues, including one hydrogen bond. Notably, these amino acids also play a crucial role in forming a hydrophobic pocket that interacts with the C-terminal helix of KAP3, and they contribute significantly to stabilizing this C-terminal helix of KAP3 through essential interactions (Figure 2c-f).

The KAP3 C-terminal helix also participates in the interaction of APC, with Gln682 and Met686 engaging APC. Notably, Met686 co-inserts into the hydrophobic pocket alongside the KIF3A 3_10_-helix, reinforcing the stability of the interface (Figure 5a-b and Supplementary Figure 8f).

In addition to these primary contacts, the β-hairpin region of KIF3A interacts with the central portion of the APC ARM domain through Asp615 and Gln620 (Figure 5b). However, binding free energy calculations suggest that the interaction involving APC residues Asn625, Thr626, and Leu627 contributes minimally to the overall binding affinity (Figure 5c).

### The Conserved Helices within the KIF3A Hitchdock Motif Dominate Specific APC Binding

Based on the functionally critical role of this H-βh-H motif in the binding of KAP3 adaptor and APC cargo as well as its structural characteristics as a protruding hook, we term it the kinesin-2 “Hitchdock domain”, as it simultaneously functions to *dock* the adaptor and *hitch* the cargo. Conservation analysis reveals that the hydrophobic core within the APC-binding pocket, where the KIF3A Hitchdock motif interacts, is highly conserved (Figure 6a). The APC interaction interface is enriched with hydrophobic residues, including multiple Leu, Ala, Ile, Val, and Met (Supplementary Figure 8f). Similarly, the KIF3A Hitchdock itself is highly conserved (Figure 2a), with the helical region contributing most of the binding energy, while the β-hairpin provides only a minor contribution (Figure 6b).

**Figure 6.**
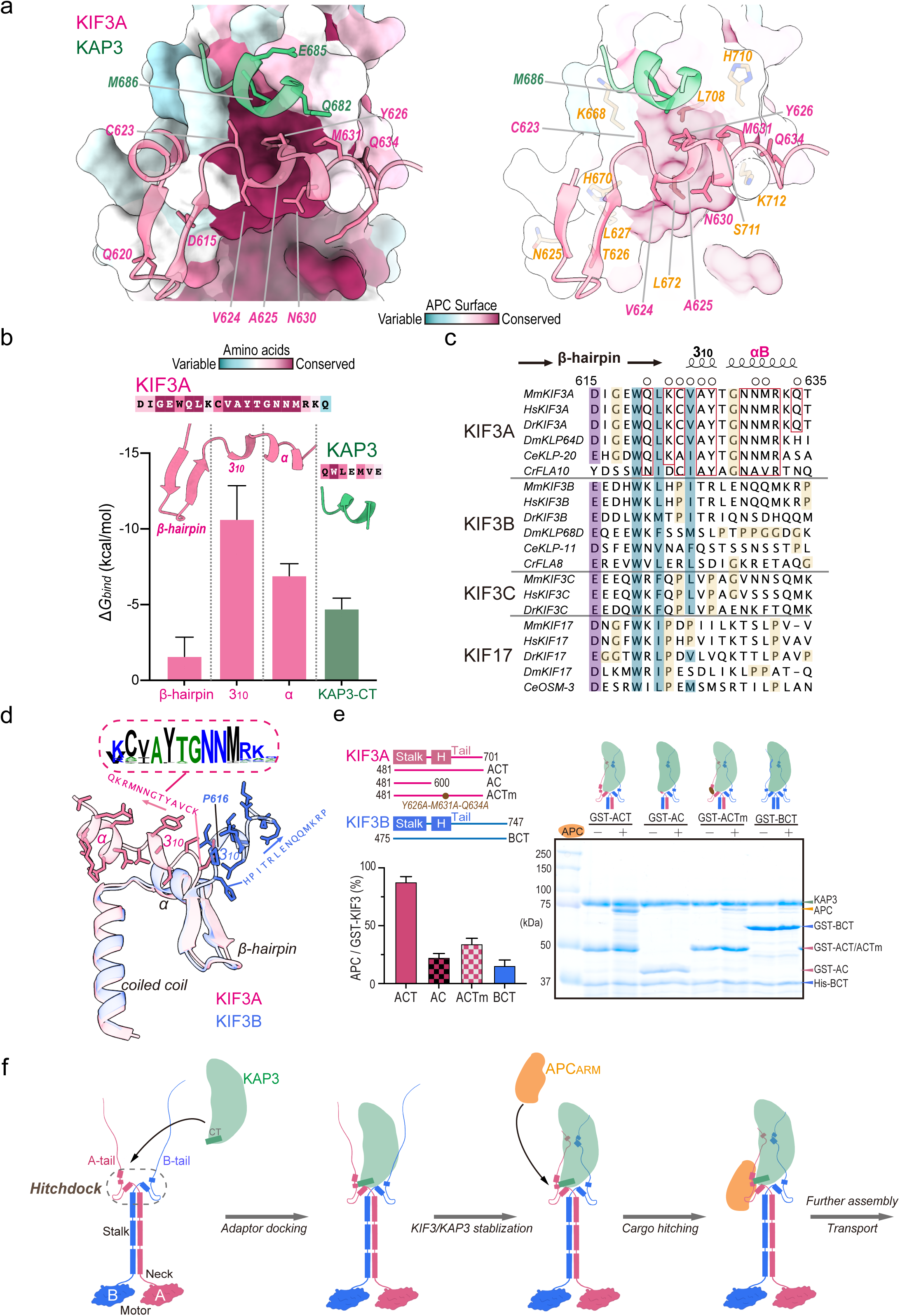
The KIF3A Hitchdock motif is critical for specific APC binding. (a) Conservation analysis of the APC binding surface. The hydrophobic core of the APC-binding pocket is highly conserved, with key interacting residues labeled. (b) Binding free energy contributions and conservation analysis of residues in the KIF3A/KAP3 interaction regions. The 3_10_-helix and α-helix contribute significantly to binding, while the β-hairpin region contributes less. (c) Multiple sequence alignment of the KIF3A Hitchdock region across kinesin-2 family members. The alignment was generated using Clustal Omega (https://www.ebi.ac.uk/jdispatcher/msa/clustalo). Purple highlights indicate conserved charged residues, cyan highlights indicate conserved hydrophobic/aromatic residues, and open circles mark residues involved in APC binding. Red boxes highlight KIF3A-specific conserved residues, and yellow highlights indicate proline residues in KIF3B/C and KIF17. (d) Structural alignment of the Hitchdock motifs in KIF3A and KIF3B. Pro616 in KIF3B, which alters structural orientation, is labeled. The conserved sequence motif in KIF3A critical for APC binding is shown (generated using WebLogo http://weblogo.berkeley.edu/). (e) Biochemical validation of the KIF3A Hitchdock motif in APC binding. GST-tagged KIF3A and KIF3B C-terminal proteins, along with KIF3A tail truncation and Hitchdock triple mutants, were used in pull-down assays with APC ARM in the presence of KIF3B C-terminal and KAP3. The left panel shows the KIF3 constructs, the middle panel summarizes the results, and the right panel shows the Coomassie-stained SDS-PAGE gel. Bar graphs indicate mean ± SD. (f) Model of KIF3 binding to adaptor and cargo First, dimerization and Hitchdock assembly: The kinesin-2 dimer is formed through stalk-mediated dimerization, assembling the Hitchdock domain in the tail. Second, adaptor docking: The KIF3A/B Hitchdock regions cooperatively recruit the KAP3 C-terminal helix, assembling the kinesin-2/adaptor complex, while the KAP3 C-terminal helix serves as an accessory component that further integrates the Hitchdock domain. Third, structural stabilization: The KIF3 tail region further stabilizes KAP3 binding, reinforcing the structural integrity of the complex. Last, cargo recognition and anchoring: The KIF3A Hitchdock motif specifically recognizes APC cargo, working in concert with KAP3 to stably anchor and transport the cargo.

Interestingly, despite a high degree of sequence similarity between KIF3B/C and KIF3A in the αA-β-hairpin region (Figure 2f), the corresponding 3_10_-αB region of KIF3B/C exhibits substantial divergence from KIF3A (Figure 6c). Within the kinesin-2β (KIF3B/C) subfamily, this region also shows relatively low conservation (Figure 2a, 6c). Notably, many KIF3B/C and KIF17 homologs contain multiple proline residues in this region, which can disrupt helical formation and potentially alter the structural conformation (Figure 6c).

Structural alignment of the Hitchdock regions in KIF3A and KIF3B reveals that the presence of Pro616 in KIF3B induces a loop reorientation following the β-hairpin and prevents the formation of the αB helix after the 3_10_-helix. This structural deviation suggests that KIF3B primarily contributes to KAP3 binding rather than APC recognition (Figure 6d). In contrast, the 3_10_-αB region of KIF3A is highly conserved, forming a C/A-V/I-A-Y-T-G-N-N-M cargo recognition motif (Figure 6d). Together with αA, this sequence motif shapes the hydrophobic pocket of the KIF3 Hitchdock fold, which interacts with the KAP3 C-terminal helix and anchors the hydrophobic interface of APC for cargo binding.

To further validate these structural findings, we conducted biochemical pull-down assays using wild-type and mutated KIF3 C-terminal protein. Based on KIF3A C-terminal constructs (ACT), we generated a KIF3A tail truncation mutant (AC) and a triple alanine substitution mutant (ACTm: Y626A-M631A-Q634A) in the 3_10_-αB region (Figure 6e). Pull-down assays showed that in the presence of KAP3 and KIF3B, wild-type ACT efficiently captured APC ARM at an approximately 1:1 ratio. However, APC binding was significantly reduced in the truncated AC mutant, the ACTm mutant, and the BCT construct lacking KIF3A (Figure 6e). These results underscore the essential role of the KIF3A Hitchdock site in APC cargo binding.

Integrating our structural, MD, and biochemical analyses, we propose the following molecular model for heterodimeric kinesin-2 adaptor and cargo recognition (Figure 6f). First, dimerization and Hitchdock assembly: The kinesin-2 dimer is formed through stalk-mediated dimerization, assembling the Hitchdock domain in the tail. Second, adaptor docking: The KIF3A/B Hitchdock regions cooperatively recruit the KAP3 C-terminal helix, assembling the Kinesin-2/Adaptor complex, while the KAP3 C-terminal helix serves as an accessory component that further integrates the Hitchdock domain; Third, structural stabilization: the KIF3 tail region further stabilizes KAP3 binding, reinforcing the structural integrity of the complex; Last, cargo recognition and anchoring: the KIF3A Hitchdock motif specifically recognizes APC cargo, working in concert with KAP3 to stably anchor and transport the cargo. This model highlights the dominant role of the KIF3A Hitchdock in coordinating adaptor and cargo specificity. While the helical regions of the KIF3A Hitchdock motif provide the primary specificity and binding affinity, the β-hairpin and KIF3B Hitchdock motif contribute structural support (Figure 6f)

## Discussion

Motor proteins, such as kinesin, dynein, and myosin, rely on adaptor proteins to ensure specific cargo recognition, transport regulation, and functional diversity. Adaptor proteins serve as bridges, linking motor proteins to organelles, vesicles, or proteins, facilitating precise identification and transport of target cargo. By interacting with various adaptor proteins, motor proteins can engage in multiple cellular processes, adapting to diverse cellular requirements ^4,5^. Our study provides the first high-resolution structural insights into the kinesin-adaptor-cargo binding, revealing a highly conserved Helix-β-hairpin-Helix motif in the KIF3 tail that serves as a central scaffold for KAP3 binding (Figures 1-2). This motif, which we term the “Hitchdock domain,” is critical for the formation of the primary binding interface, including between KIF3 and KAP3 (ABK interface1), which stabilize the KIF3/KAP3 interaction through hydrophobic pockets, electrostatic interactions, and hydrogen bonding networks (Figure 2). Notably, KIF3A plays a dominant role in APC binding, with its 3_10_-αB region engaging a unique hydrophobic pocket of the ARM domain of APC (Figures 5-6). In contrast, KIF3B contributes primarily to stabilizing the KIF3/KAP3 complex through additional interfaces (ABK interface2, BK interfaces 1-2) (Figure 3) but does not directly participate in APC recognition (Figures 4-6). This functional divergence is likely due to structural constraints in KIF3B, such as the presence of proline residues that disrupt helical formation in its 3_10_-αB region (Figure 6). These findings underscore the importance of the Hitchdock region in kinesin-2 cargo recognition (Figure 6f) and provide a structural basis for understanding how KIF3A and KAP3 cooperate to achieve specific cargo transport.

The discovery of the hook-like kinesin-2 Hitchdock-KAP3 structure extends beyond kinesin-2, suggesting parallels and a conserved mechanism among other molecular motors, such as kinesin-1 and dynein, which also utilize conserved hook-like structures for cargo recognition (Supplementary Figure 9). Despite differences in composition, the recently resolved structure of the cytoplasmic dynein FTS–HOOK– FHIP1B (FHF) adaptor complex exhibits a strikingly similar hook-like architecture to our KIF3 tail/KAP3 structure ^33^. The dynein/dynactin-associated HOOK protein contains a long coiled-coil stalk-like region, analogous to the stalk of kinesins, with its C-terminal coiled-coil and extension (CTE) helices mediating interactions with both FHIPB and FTS ^32,33,36^. This interaction forms a complex highly reminiscent of the KIF3 tail/KAP3 structure analyzed in our study, probably serving as a tethering center for adaptors and cargo in a manner similar to the Hitchdock region in kinesin-2 (Supplementary Figure 9a). Moreover, the cargo-binding region formed by the C-terminal region of the kinesin-1 heavy chain and its light chain (KLC) is also proposed to adopt a hook-like structure ^8^. Interestingly, the tetratricopeptide repeat (TPR) domain of KLC shares structural similarity with the ARM domain of KAP3, suggesting that their overall architecture may resemble the Hitchdock-KAP3 structure we resolved (Supplementary Figure 9a), though this awaits further validation.

Additionally, kinesin-1 cargo employs tyrosine-acidic and tryptophan-acidic motifs for binding with the conserved region of KLC ^10–12^, while dynein cargos also rely on CC box, HBS1 and spindly motifs for binding with the conserved region of dynein or adaptors ^37,38^. This suggests a shared mechanistic framework among molecular motors, where conserved structural motifs enable specific cargo recognition. Taken together, we speculate that other motor proteins may also form tethering centers, akin to the Hitchdock region, at their tail ends or through primary adaptors, subsequently adopting hook-like architectures to facilitate coordinated interactions with adaptors and cargo. We also propose a unified model for cargo recognition in kinesin and dynein. The C-terminus of adaptors in kinesin 1/2 and dynein features a helix-based adaptor/cargo tethering site that promotes adaptor assembly and facilitates subsequent cargo recognition and binding (Supplementary Figure 9b). The role of the tethering site in cargo binding may vary: for kinesin-1 and dynein, cargo specificity arises from the interaction between cargo and adaptor proteins, while in kinesin-2, cargo specificity is determined by the tethering site itself.

Furthermore, the Hitchdock region contains phosphorylatable tyrosine residues (e.g., Tyr626 ^39^), which may serve as regulatory sites to modulate KIF3 binding to APC. Such post-translational modifications could integrate kinesin-2 function into broader cellular signaling networks, such as the Wnt pathway ^40^, highlighting the potential for dynamic regulation of intracellular transport processes.

Our findings also have significant theoretical and practical implications by the discovery of the conserved cargo recognition motif (C/A-V/I-A-Y-T-G-N-N-M) in the KIF3 Hitchdock region, which could promote further exploration of specific cargos, expanding our understanding of the kinesin-2-mediated transport mechanism as well as the pathogenesis of kinesin-2-related diseases. By elucidating the structural basis of KIF3/KAP3/APC interactions, this study advances our understanding of kinesin-2 function and its role in cellular transport. The Hitchdock region represents a potential therapeutic target for diseases linked to kinesin-2 dysfunction, such as ciliopathies and cancer. However, certain limitations remain. For example, the high flexibility of some regions in the KIF3 tail and KAP3 prevented their high-resolution analysis in our cryo-EM maps (Figure 1e), leaving open the possibility of additional binding or regulatory sites. Future studies could employ advanced structural techniques or biochemical assays to explore these unresolved regions and investigate the dynamic regulatory mechanisms of the Hitchdock region, such as phosphorylation-dependent modulation of cargo binding.

In conclusion, this study delineates the structural and functional architecture of the KIF3/KAP3 complex, emphasizing the critical role of the Hitchdock region in cargo recognition and transport. These findings not only deepen our understanding of kinesin-2 biology but also provide a framework for exploring shared mechanisms among molecular motors. Future research should focus on the regulatory potential of the Hitchdock region and its integration into cellular signaling networks, offering new avenues for therapeutic intervention in kinesin-2-related diseases.

## Materials and Methods

### Constructs, Protein Expression and Purification

The gene encoding the C-terminal region of *Mus musculus* KIF3A (NCBI accession: NP_032469.2; residues 481–701, ACT) was cloned into a pETDuet-1 vector with an N-terminal 6His tag, while the C-terminal region of *M. musculus* KIF3B (NCBI accession: NP_032470.3; residues 475–747, BCT) inserted into another multiple cloning site (MCS) without expression tags for the co-expression of the KIF3A/B tail heterodimer. These sequences were also cloned into a pGEX-6p-3 vector containing an N-terminal GST tag for pull-down assays. A truncated form of KIF3A (AC, residues 481–600) and a triple point-mutant variant (ACTm) were generated using PCR and cloned into the pGEX-6p-3 vector.

*M. musculus* KAP3A (residues 1–693, NCBI accession: NP_001292572.1) and the ARM domain of *M. musculus* APC (NCBI accession: AAB59632.1; residues 338– 1010) were PCR-amplified and cloned into a pET-21b vector without expression tags. Recombinant plasmids were transformed into Escherichia coli BL21 (DE3) (Novagen), and both KIF3A/B/KAP3 and KIF3A/B/KAP3-APC complexes were reconstituted and purified as previously described ^9^. Purified proteins were concentrated to 10 mg/mL using ultrafiltration, flash-frozen in liquid nitrogen, and stored at −80°C.

### Cryo-EM Sample Preparation and Data Collection

For cryo-EM analysis of both KIF3 C-terminus/KAP3 and KIF3 C-terminus/KAP3/APC_ARM_ complexes, 3 μL of the sample at a concentration of 1.5 mg/mL was applied onto a hydrophilized holey carbon grid (Cu, R1.2/1.3, 300 mesh, Quantifoil). The grid was then blotted for 4 s (blot force 10) and plunge-frozen in liquid ethane using a Vitrobot Mark IV (FEI) at 6°C and 100% humidity.

Images were acquired on a CRYO ARM^TM^ 200 (JEOL) microscope operating at 200 kV using SerialEM for automated data collection. Movie frames were recorded with a Gatan K3 direct electron detector operated in counting mode and correlated-double sampling mode at a nominal magnification of 80,000×, yielding a pixel size of 0.571 Å/pixel. The data were collected in zero-loss mode with an energy filter slit width of 20[eV and an objective lens aperture diameter of 150 μm. For the KIF3 C-terminus/KAP3 sample, 34,176 movies were recorded with a total electron exposure of 65.3 e[/Å² over 75 frames. For the KIF3 C-terminus/KAP3-APC_ARM_ sample, 36,020 movies were recorded with a total electron exposure of 64.7 e[/Å² over 80 frames. The defocus range was set from −0.6 to −1.8 μm in 0.1 μm increments. A summary of the imaging parameters is provided in Supplementary Table 1.

### Cryo-EM Single-particle Data Processing

All data was processed using CryoSPARC (version 4.6.2) ^41^. First, raw movies were subjected to Patch Motion Correction to compensate for beam-induced sample drift, followed by Patch CTF Estimation to determine contrast transfer function (CTF) parameters. Dose-weighted images with a CTF fit resolution worse than 8 Å were discarded, leaving 33,576 and 35,534 images for the KIF3 tail/KAP3 and KIF3 tail/KAP3-APC_ARM_ datasets, respectively, for further particle picking. For particle selection, an initial Blob Picker job was used to generate preliminary 2D templates. These templates were refined through 2D classification, and the improved templates were then used for Template Picker to enhance the accuracy of particle selection. To mitigate orientation bias, multiple rounds of template picking were conducted with varying particle sizes to ensure diverse angular coverage. Extracted particles were down-sampled to a pixel size of 2.284 Å (4× binning) with a box size of 228 Å for initial processing. Duplicates from multiple rounds of template picking were merged and removed. Particles were classified into 300 classes, and poorly defined classes were discarded. Several rounds of 2D classification were performed using different mask sizes (90 Å, 110 Å, 130 Å, 150 Å, and 170 Å) to capture particles with a broader range of orientations. Well-defined 2D classes from these classifications were selected, merged, and deduplicated. This resulted in 9,701,817 particles for KIF3 tail/KAP3 and 6,456,283 particles for KIF3 tail/KAP3-APC_ARM_, which were used for 3D reconstruction.

A subset of particles was used to generate six initial 3D models using Ab-initio Reconstruction. These models were then employed in Heterogeneous Refinement to classify particles into distinct conformational or compositional states. To improve data quality, selected 3D classes were re-extracted at a pixel size of 1.142 Å (2× binning) and underwent multiple rounds of Ab-initio Reconstruction and Heterogeneous Refinement to discard low-quality particles. For KIF3 tail/KAP3, two well-defined classes with good orientation distributions were selected and refined separately using Non-Uniform Refinement ^42^ to improve map quality and resolution.

In contrast, KIF3 tail/KAP3-APC exhibited significant orientation bias. To address this, the best two classes were combined and subjected to Non-Uniform Refinement, followed by orientation rebalance, where dominant orientations were randomly down-sampled to reduce bias. Focused refinement was applied to regions of interest to enhance local structural details using Local Refinement. Moreover, reference-based motion correction and global/local CTF refinement were performed to further improve the final reconstruction quality and resolution, and local resolution estimation was conducted to assess the map quality. The final map was sharpened using DeepEMhancer ^43^. A summary of the data processing workflow and final reconstruction resolutions is provided in Supplementary Figure 1 for KIF3 tail/KAP3 and Supplementary Figure 5 for KIF3 tail/KAP3-APC_ARM_.

### Model Building and Validation

For KIF3 tail/KAP3, the Class 1 sharpened map was used for de novo model building with ModelAngelo ^34^ to generate an initial model. The raw map was used to supplement the construction of the linking loop in the KIF3A tail region. The model was then imported into Phenix ^44^ for automated real-space refinement, followed by manual corrections using Coot ^45^.

For the KIF3 tail/KAP3-APC, the initial model was constructed by docking the KIF3 tail/KAP3 model solved in this study and integrating the available crystal structure of the APC ARM domain complex (PDB: 4YJE). The model was further refined using MD-based flexible fitting with Cryo_fit in Phenix, followed by manual adjustments in Coot to improve alignment with the EM density. The overall model-building workflow is illustrated in Supplementary Figure 2 (for KIF3 tail/KAP3) and Supplementary Figure 5 (for KIF3 tail/KAP3-APC_ARM_).

Model validation was performed using MolProbity ^46^ to assess stereochemical quality. A summary of model-building and refinement statistics is provided in Supplementary Table 1.

### Molecular Dynamic Simulation and Free Energy Calculation

The MD simulations of KIF3 tail/KAP3 and KIF3 tail/KAP3-APC were performed using GROMACS (version 2024.4), with slight modifications to the protocol previously described ^47^. Briefly, the topology files were generated using the OPLS-AA force field parameter set. Each system was solvated in 150 mM NaCl with SPC/E water models in a cubic box. Neutralizing counterions were added, and the steepest descent energy minimization was performed. This was followed by a two-step equilibration: 100 ps of isochoric–isothermal equilibration (NVT) and 100 ps of isothermal–isobaric equilibration (NPT). All position restraints were removed, and simulations were run for 1 ns. Root mean square deviation (RMSD) analyses were used to calculate the standard deviation (SD) of atomic positions for specified amino acids, compared to their initial positions within the energy-minimized and equilibrated structures. Each simulation was performed in triplicate.

Based on the RMSD stability observed during the 1 ns MD simulation, we selected 20 stable frames from the 0.8 to 1 ns interval to estimate the binding free energy between the proteins using the gmx_MMPBSA tool ^48^. The molecular mechanics/Poisson-Boltzmann surface area (MM/PBSA) method was applied, partitioning the free energy into molecular mechanical energies (electrostatic and van der Waals), polar solvation energies (calculated via the Poisson-Boltzmann equation), and non-polar solvation energies (estimated from the solvent-accessible surface area). To identify key residues contributing to binding, per-residue energy decomposition was performed using the decomposition feature of gmx_MMPBSA. The data of total delta free energy were plotted using GraphPad Prism7 (RRID: SCR_002798).

### Crosslinking Mass Spectrometry

To investigate potential interactions among the KIF3A/B tail, KAP3, and APC, the KIF3 tail/KAP3-APC_ARM_ complex was crosslinked using bis(sulfosuccinimidyl) suberate (BS3; Dojindo) for 10 minutes at room temperature, followed by quenching with 50 mM Tris-HCl. The sample was then digested with trypsin, desalted, and analyzed using a ZenoTOF7600 mass spectrometer (SCIEX) coupled with an UltiMate3000 RSLCnano system (Thermo Fisher). Crosslinked fragments containing BS3 were analyzed using MaxLynx within the MaxQuant software suite ^49^. A summary of detected crosslinks is provided in Supplementary Table 2.

### Conservation Analysis

To evaluate the evolutionary conservation of the proteins KIF3A, KIF3B, KAP3, and APC within our structural models, we utilized the ConSurf server ^50^ to analyze the phylogenetic relationships among homologous sequences to determine the conservation levels of amino acid positions. The amino acid sequences of KIF3A, KIF3B, KAP3, and APC were submitted to ConSurf, which automatically identified homologous sequences using the HMMER algorithm against the UniRef90 database, applying an E-value cutoff of 0.0001 to ensure significant matches. Redundant sequences were filtered out, and the remaining sequences were aligned using the MAFFT-L-INS-i algorithm for multiple sequence alignment (MSA). A phylogenetic tree was constructed from the MSA using the neighbor-joining method. Subsequently, the Rate4Site algorithm calculated evolutionary conservation scores for each amino acid position, assigning grades from 1 (most variable) to 9 (most conserved) based on evolutionary rates. These conservation scores were then mapped onto the three-dimensional structures of the proteins, facilitating visualization of conserved and variable regions.

### Sequence and Structural Analysis

Structure-based multiple sequence alignments were performed using Clustal Omega (RRID: SCR_001591) and ESPript server (RRID: SCR_006587). The protein sequences used for analysis were obtained via the NCBI protein database under the accession numbers NP_001287720.1 (*Homo sapiens* KIF3A), NP_001277734.1 (*M. musculus* KIF3A), NP_001093615.1, (*M. musculus* KIF3B), NP_001093615.1, NP_001093615.1 (*Danio rerio* KIF3A), (*Danio rerio* KIF3B), NP_523934.1 (*Drosophila* Klp64D), NP_497178.1 (*C. elegans* Klp-20), XP_001701510.1 (*Chlamydomonas* FLA10), NP_004789.1 (*H. sapiens* KIF3B), NP_032470.3 (*M. musculus* KIF3B), NP_001093615.1 (*D. rerio* KIF3B), NP_524029.2 (*Drosophila* Klp64D), NP_741473.1 (*C. elegans* Klp-11), and XP_001697037.1 (*Chlamydomonas* FLA8), NP_002245.4 (*Homo sapiens* KIF3C), NP_032471.2 (*M. musculus* KIF3C), XP_002661420.3 (*Danio rerio* KIF3C), NP_065867.2 (*Homo sapiens* KIF17), NP_034753.1 (*M. musculus* KIF17), XP_068080281.1 (*Danio rerio* KIF17), NP_651939.4 (*Drosophila* KIF17), NP_001367796.1 (*C. elegans* OSM-3), NP_001191443.1 (*Homo sapiens* KAP3), NP_035288.2 (*M. musculus* KAP3), NP_001004644.1 (*Danio rerio* KAP3), NP_001138186.1 (*Drosophila* KAP3), NP_001021247.1 (*C. elegans* KAP-1), XP_001698323.1 (*Chlamydomonas* KAP), NP_000029.2 (*Homo sapiens* APC) NP_001389660.1 (*M. musculus* APC), NP_001137312.1 (*Danio rerio* APC), NP_001263046.1 (*Drosophila* APC). Structural analysis was performed, and structural figures were generated using the program UCSF ChimeraX (RRID: SCR_015872).

### Pull-down Assay

For the pull-down analysis using wild-type KIF3A/B tail and mutated KIF3A tail, E. coli cells expressing the bait proteins (GST-fused C-terminal KIF3A or KIF3B constructs) (Fig. 6e) were premixed with E. coli cells overexpressing input proteins (KIF3B tail, KAP3, and APC_ARM_). The mixtures were lysed by sonication in buffer containing 20 mM Tris-HCl (pH 8.0), 150 mM NaCl, 7 mM β-mercaptoethanol, and 5% (v/v) glycerol.

After centrifugation, the cleared supernatants were incubated with pre-equilibrated Glutathione Sepharose™ 4B beads (Cytiva) for 1 hour at 4°C on a rocker. The bound proteins were eluted with 10 mM reduced glutathione and analyzed by SDS-PAGE followed by Coomassie Brilliant Blue (CBB) staining. All experiments were performed in triplicate. Quantification was performed using ImageJ (RRID: SCR_003070), and statistical analysis was conducted with GraphPad Prism7 (RRID: SCR_002798).

## Acknowledgments

We thank the current members of the MK and NH laboratory for their valuable assistance. This research was supported by JST ERATO Grant Number JPMJER2202 (MK), and the JSPS KAKENHI Grant Numbers JP23000013 (NH), JP16H06372 (NH), JP21H05247 (MK), JP24K18106 (XJ), JP22H02554 (RD) and JP24KF0141 (MK & XJ). This research was partially supported by the Platform Project for Supporting Drug Discovery and Life Science Research (Basis for Supporting Innovative Drug Discovery and Life Science Research (BINDS)) from AMED under Grant Numbers JP23ama121002 and JP23ama121018. In addition, XJ acknowledges the support from JSPS as a JSPS international research fellow.

## Author contributions

Conceptualization: NH, MK; Methodology: XJ, RD, SO, MK; Investigation: XJ; Data curation: XJ, RD, HY, BN, SO; Funding acquisition: MK, NH, XJ, RD; Project administration: NH, MK; Validation: NH, MK, RD; Supervision: NH, MK; Writing – original draft: XJ; Writing – review & editing: XJ, RD, MK, NH.

## Data and materials availability

Structural data will be deposited to the corresponding database. Other data are available from the corresponding author on request.

## Supporting Information

Supplementary Figures 1-9, Supplementary Tables 1-2

## Supplementary Figures

**Supplementary Figure 1.** Cryo-EM data processing workflow for the KIF3A/B/KAP3 complex.

**Supplementary Figure 2.** Model building for the KIF3A/B/KAP3 complex.

**Supplementary Figure 3.** KIF3A/B interaction and the binding contribution of KAP3 in ABK interface2.

(a) Ribbon representation of the KIF3A/B stalk-tail junction. Key interacting residues and hydrogen bonds are labeled.

(b) Cartoon representation of the ABK interface2 structure and hydrogen bond network (left). Binding free energy contributions of KAP3 residues involved in the interface are summarized (right). Key interacting residues are labeled.

(c) Multiple sequence alignment of KAP3 residues involved in ABK interface2 across species. Key interacting residues are labeled.

**Supplementary Figure 4.** Surface hydrophobicity and charge of KIF3A/B and KAP3.

(a, b) Hydrophobic and electrostatic properties of the KAP3 surface, with KIF3 structures shown as cartoons.

(c) Hydrophobic and electrostatic properties of the KIF3 surface, with KAP3 structures shown as cartoons.

**Supplementary Figure 5.** Cryo-EM data processing workflow for the KIF3A/B/KAP3-APC complex.

**Supplementary Figure 6.** Model building for the KIF3A/B/KAP3-APC complex.

**Supplementary Figure 7.** Crosslinking mass spectrometry (XL-MS) results of the KIF3A/B/KAP3-APC complex.

(a) Circular representation (left) and 2D diagram (right) of XL-MS results for the KIF3 C-terminal/KAP3-APC complex on the constructs. Intermolecular crosslinks are shown in purple, and intramolecular crosslinks are shown in gray. Only crosslinks visible in the structural model are displayed. Full data are available in Supplementary Figure 7 and Supplementary Table 2.

(b) XL-MS results mapped onto the structural model. Intermolecular crosslinks are depicted using dashed lines in two colors, each corresponding to the respective components, while intramolecular crosslinks are represented by dashed lines in a single color matching the component. Crosslink pairs are within theoretical distances.

**Supplementary Figure 8.** Molecular dynamics and MMPBSA analysis of the KIF3A/B/KAP3-APC complex.

(a) RMSD changes during three replicate MD simulations.

(b–e) RMSD changes of APC, KAP3, KIF3A, and KIF3B residues before and after MD simulations.

(f) Multiple sequence alignment of KAP3 and APC residues involved in KIF3/KAP3-APC binding across species. Key interacting residues are labeled. The alignment was generated using Clustal Omega (https://www.ebi.ac.uk/jdispatcher/msa/clustalo).

**Supplementary Figure 9.** Hook-like cargo-binding structures suggest a shared mechanism in kinesin and dynein cargo/adaptor assembly.

(a) Comparison of the KIF3/KAP3 cargo-binding module map obtained in this study with the dynein FTS-HOOK-FHIP1B (FTF) cargo-binding adaptor complex map and a possible kinesin-1 KHC/KLC model. The FTF density is derived from (EMDB: EMD-18302), and the density of the KLC1 TPR domain is generated based on its crystal structure (PDB: 3NF1).

(b) A proposed common model for kinesin/dynein cargo recognition mechanism. The C-terminus of kinesin 1/2 and dynein adaptors contains an adaptor/cargo tethering site formed by helices, which mediates adaptor assembly and further recognition and binding to cargo. The form and contribution of the tethering site in cargo binding may differ; for kinesin-1 and dynein, cargo specificity is provided by the cargo-adaptor protein interaction (dashed line), while for kinesin-2, it is provided by the tethering site itself.

**Supplementary Table 1.** Cryo-EM data collection, modeling, and refinement statistics.

**Supplementary Table 2.** Cross-links detected by cross-linking mass spectrometry.

## References

1. Hirokawa, N. Kinesin and dynein superfamily proteins and the mechanism of organelle transport. Science (1979) 279, 519–26 (1998).

2. Vale, R. D. The molecular motor toolbox for intracellular transport. Cell 112, 467–480 (2003).

3. Hirokawa, N., Niwa, S. & Tanaka, Y. Molecular motors in neurons: Transport mechanisms and roles in brain function, development, and disease. Neuron 68, 610– 638 (2010).

4. Ahmet Yildiz. Mechanism and regulation of kinesin motors. Nat Rev Mol Cell Biol 26, 86–103 (2025).

5. Hirokawa, N., Noda, Y., Tanaka, Y. & Niwa, S. Kinesin superfamily motor proteins and intracellular transport. Nat Rev Mol Cell Biol 10, 682–696 (2009).

6. Hirokawa, N. & Tanaka, Y. Kinesin superfamily proteins (KIFs): Various functions and their relevance for important phenomena in life and diseases. Exp Cell Res 334, 16– 25 (2015).

7. Lee Sweeney, H. & Holzbaur, E. L. F. Motor Proteins. Cold Spring Harb Perspect Biol 10, a021931 (2018).

8. Tan, Z. et al. Autoinhibited kinesin-1 adopts a hierarchical folding pattern. Elife 12, (2023).

9. Jiang, X. et al. The two-step cargo recognition mechanism of heterotrimeric kinesin. EMBO Rep 24, e56864 (2023).

10. Pernigo, S., Lamprecht, A., Steiner, R. A. & Dodding, M. A. Structural Basis for Kinesin-1:Cargo Recognition. Science (1979) 340, 356–360 (2013).

11. Pernigo, S. et al. Structural basis for isoform-specific kinesin-1 recognition of Y-acidic cargo adaptors. Elife 7, e38362 (2018).

12. Quyen Nguyen, T., et al. Characterization of the binding mode of JNK-interacting protein 1 (JIP1) to kinesin-light chain 1 (KLC1). Journal of Biological Chemistry 293, (2018).

13. Takeda, S. et al. Kinesin superfamily protein 3 (KIF3) motor transports fodrin-associating vesicles important for neurite building. Journal of Cell Biology (2000) doi:10.1083/jcb.148.6.1255.

14. Scholey, J. M. Kinesin-II, a membrane traffic motor in axons, axonemes, and spindles. Journal of Cell Biology 133, 1–4 (1996).

15. Scholey, J. M. Kinesin-2: A family of heterotrimeric and homodimeric motors with diverse intracellular transport functions. Annu Rev Cell Dev Biol 29, 443–469 (2013).

16. Kondo, S. et al. KIF3A is a new microtubule-based anterograde motor in the nerve axon. Journal of Cell Biology 125, 1095–1107 (1994).

17. Yamazaki, H., Nakata, T., Okada, Y. & Hirokawa, N. KIF3A/B: A heterodimeric kinesin superfamily protein that works as a microtubule plus end-directed motor for membrane organelle transport. Journal of Cell Biology 130, 1387–1399 (1995).

18. Yamazaki, H., Nakata, T., Okada, Y. & Hirokawa, N. Cloning and characterization of KAP3: A novel kinesin superfamily-associated protein of KIF3A/3B. Proc Natl Acad Sci U S A 93, 8443–8448 (1996).

19. Rosenbaum, J. L. & Witman, G. B. Intraflagellar transport. Nat Rev Mol Cell Biol 3, 813–825 (2002).

20. Hirokawa, N., Tanaka, Y. & Okada, Y. Cilia, KIF3 molecular motor and nodal flow. Curr Opin Cell Biol 24, 31–39 (2012).

21. Teng, J. et al. The KIF3 motor transports N-cadherin and organizes the developing neuroepithelium. Nat Cell Biol 7, 474–482 (2005).

22. Lolkema, M. P. et al. The von Hippel-Lindau tumour suppressor interacts with microtubules through kinesin-2. FEBS Lett 581, 4571–4576 (2007).

23. Carpenter, B. S., Barry, R. L., Verhey, K. J. & Allen, B. L. The heterotrimeric kinesin-2 complex interacts with and regulates GLI protein function. J Cell Sci 128, 1034–1050 (2015).

24. Ichinose, S., Ogawa, T. & Hirokawa, N. Mechanism of Activity-Dependent Cargo Loading via the Phosphorylation of KIF3A by PKA and CaMKIIa. Neuron 87, 1022– 1035 (2015).

25. Ichinose, S., Ogawa, T., Jiang, X. & Hirokawa, N. The Spatiotemporal Construction of the Axon Initial Segment via KIF3/KAP3/TRIM46 Transport under MARK2 Signaling. Cell Rep 28, 2413–2426.e7 (2019).

26. Yoshihara, S. et al. Betaine ameliorates schizophrenic traits by functionally compensating for KIF3-based CRMP2 transport. Cell Rep 35, 108971 (2021).

27. Wang, S., Tanaka, Y., Xu, Y., Takeda, S. & Hirokawa, N. KIF3B promotes a PI3K signaling gradient causing changes in a Shh protein gradient and suppressing polydactyly in mice. Dev Cell 57, 2273–2289.e11 (2022).

28. Alsabban, A. H., Morikawa, M., Tanaka, Y., Takei, Y. & Hirokawa, N. Kinesin Kif3b mutation reduces NMDAR subunit NR 2A trafficking and causes schizophrenia-like phenotypes in mice. EMBO J 39, e101090 (2020).

29. Jimbo, T. et al. Identification of a link between the tumour suppressor APC and the kinesin superfamily. Nat Cell Biol 4, 323–327 (2002).

30. Baumann, S. et al. A reconstituted mammalian APC-kinesin complex selectively transports defined packages of axonal mRNAs. Sci Adv 6, eaaz1588 (2020).

31. Garbouchian, A., Montgomery, A., Gilbert, S. P. & Bentley, M. KAP is the neuronal organelle adaptor for Kinesin-2 KIF3AB and KIF3AC. Mol Biol Cell 33, ar133 (2022).

32. Xu, L. et al. An FTS/Hook/p107 Complex Interacts with and Promotes Endosomal Clustering by the Homotypic Vacuolar Protein Sorting Complex. Mol Biol Cell 19, 5059–5071 (2008).

33. Abid Ali, F., et al. KIF1C activates and extends dynein movement through the FHF cargo adapter. Nat Struct Mol Biol (2025) doi:10.1038/s41594-024-01418-z.

34. Jamali, K. et al. Automated model building and protein identification in cryo-EM maps. Nature 628, 450 (2024).

35. Preitner, N. et al. APC is an RNA-binding protein, and its interactome provides a link to neural development and microtubule assembly. Cell 158, (2014).

36. Kendrick, A. A. et al. Hook3 is a scaffold for the opposite-polarity microtubule-based motors cytoplasmic dynein-1 and KIF1C. Journal of Cell Biology 218, 2982–3001 (2019).

37. Chaaban, S. & Carter, A. P. Structure of dynein–dynactin on microtubules shows tandem adaptor binding. Nature 610, 212–216 (2022).

38. Reck-Peterson, S. L., Redwine, W. B., Vale, R. D. & Carter, A. P. The cytoplasmic dynein transport machinery and its many cargoes. Nat Rev Mol Cell Biol 19, 382–398 (2018).

39. Hornbeck, P. V., et al. PhosphoSitePlus, 2014: Mutations, PTMs and recalibrations. Nucleic Acids Res 43, D512–D520 (2015).

40. Votin, V., Nelson, W. J. & Barth, A. I. M. Neurite outgrowth involves adenomatous polyposis coli protein and β-catenin. J Cell Sci 118, 5699–5708 (2005).

41. Punjani, A., Rubinstein, J. L., Fleet, D. J. & Brubaker, M. A. CryoSPARC: Algorithms for rapid unsupervised cryo-EM structure determination. Nat Methods 14, 290–296 (2017).

42. Punjani, A., Zhang, H. & Fleet, D. J. Non-uniform refinement: adaptive regularization improves single-particle cryo-EM reconstruction. Nat Methods 17, 1214–1221 (2020).

43. Sanchez-Garcia, R., et al. DeepEMhancer: a deep learning solution for cryo-EM volume post-processing. Commun Biol 4, (2021).

44. Liebschner, D. et al. Macromolecular structure determination using X-rays, neutrons and electrons: Recent developments in Phenix. Acta Crystallogr D Struct Biol (2019) doi:10.1107/S2059798319011471.

45. Emsley, P. & Cowtan, K. Coot: Model-building tools for molecular graphics. Acta Crystallogr D Biol Crystallogr (2004) doi:10.1107/S0907444904019158.

46. Williams, C. J. et al. MolProbity: More and better reference data for improved all-atom structure validation. Protein Science 27, 293–315 (2018).

47. Jiang, X. et al. DYF-5/MAK–dependent phosphorylation promotes ciliary tubulin unloading. Proceedings of the National Academy of Sciences 119, e2207134119 (2022).

48. Valdés-Tresanco, M. S., Valdés-Tresanco, M. E., Valiente, P. A. & Moreno, E. Gmx_MMPBSA: A New Tool to Perform End-State Free Energy Calculations with GROMACS. J Chem Theory Comput 17, 6281–6291 (2021).

49. Yllmaz, Ş., Busch, F., Nagaraj, N. & Cox, J. Accurate and Automated High-Coverage Identification of Chemically Cross-Linked Peptides with MaxLynx. Anal Chem 94, 1608–1617 (2022).

50. Yariv, B. et al. Using evolutionary data to make sense of macromolecules with a “face-lifted” ConSurf. Protein Science 32, e4582 (2023).

