## Supplemental figures and tables for "The Hitchdock Domain in Kinesin-2 Tail Enables Adaptor Assembly and Cargo Binding"

**The file includes:**

Figures S1 to S9
Tables S1 to S2

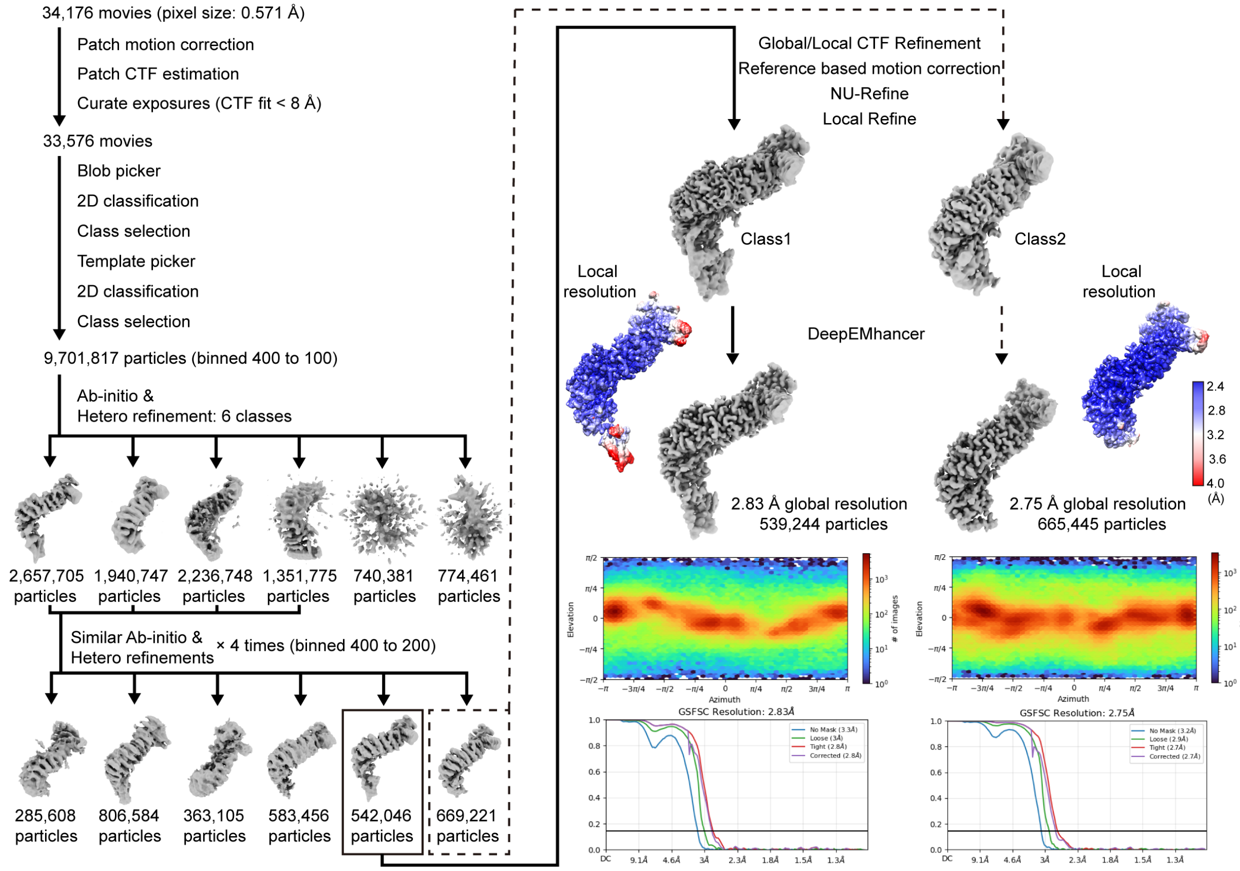


Supplementary Figure 1. Cryo-EM data processing for the KIF3A/B/KAP3 complex

**Fig. S1. Cryo-EM data processing workflow for the KIF3A/B/KAP3 complex.**

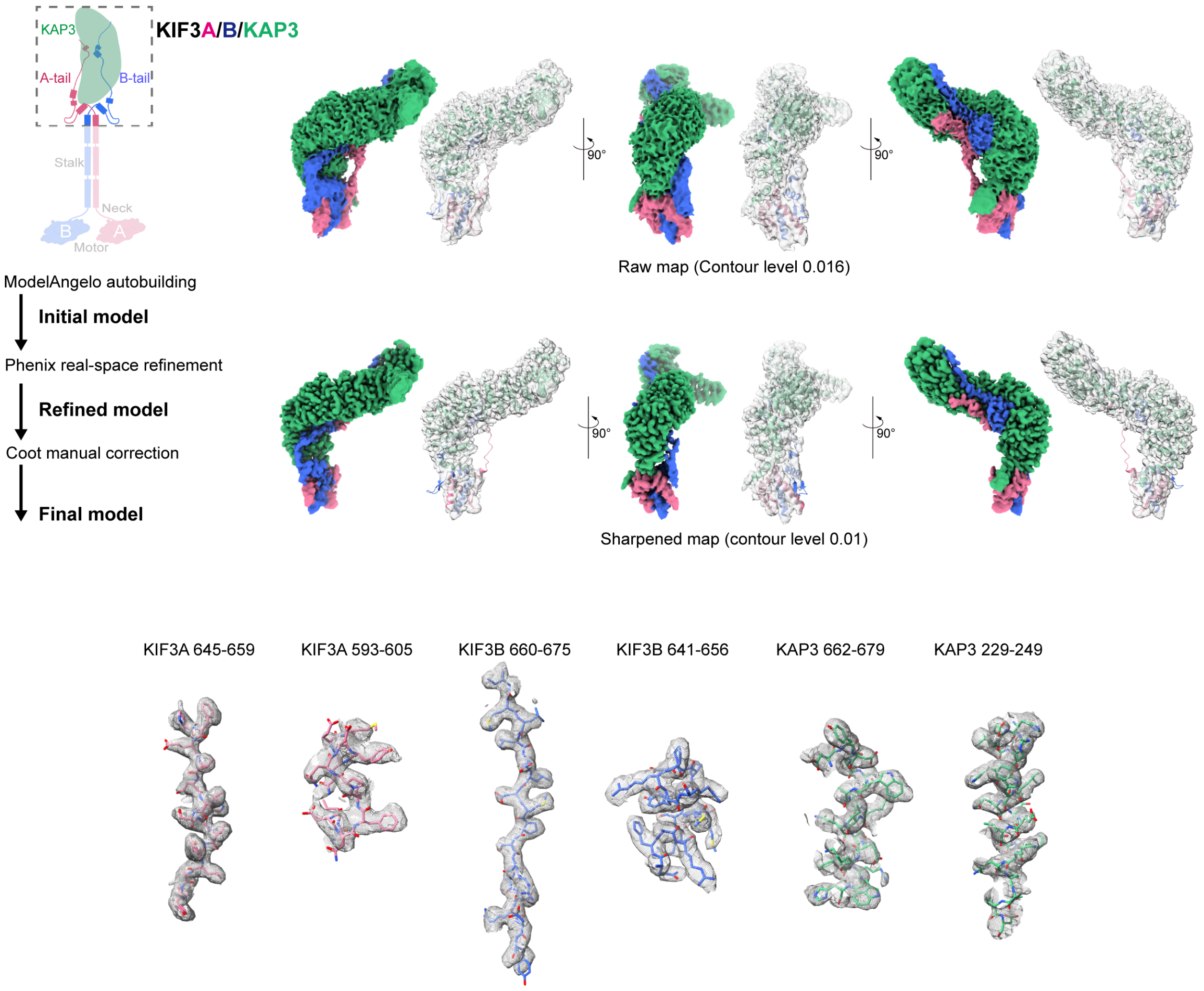


Supplementary Figure 2. Model building for the KIF3A/B/KAP3 complex

**Fig. S2. Model building for the KIF3A/B/KAP3 complex.**

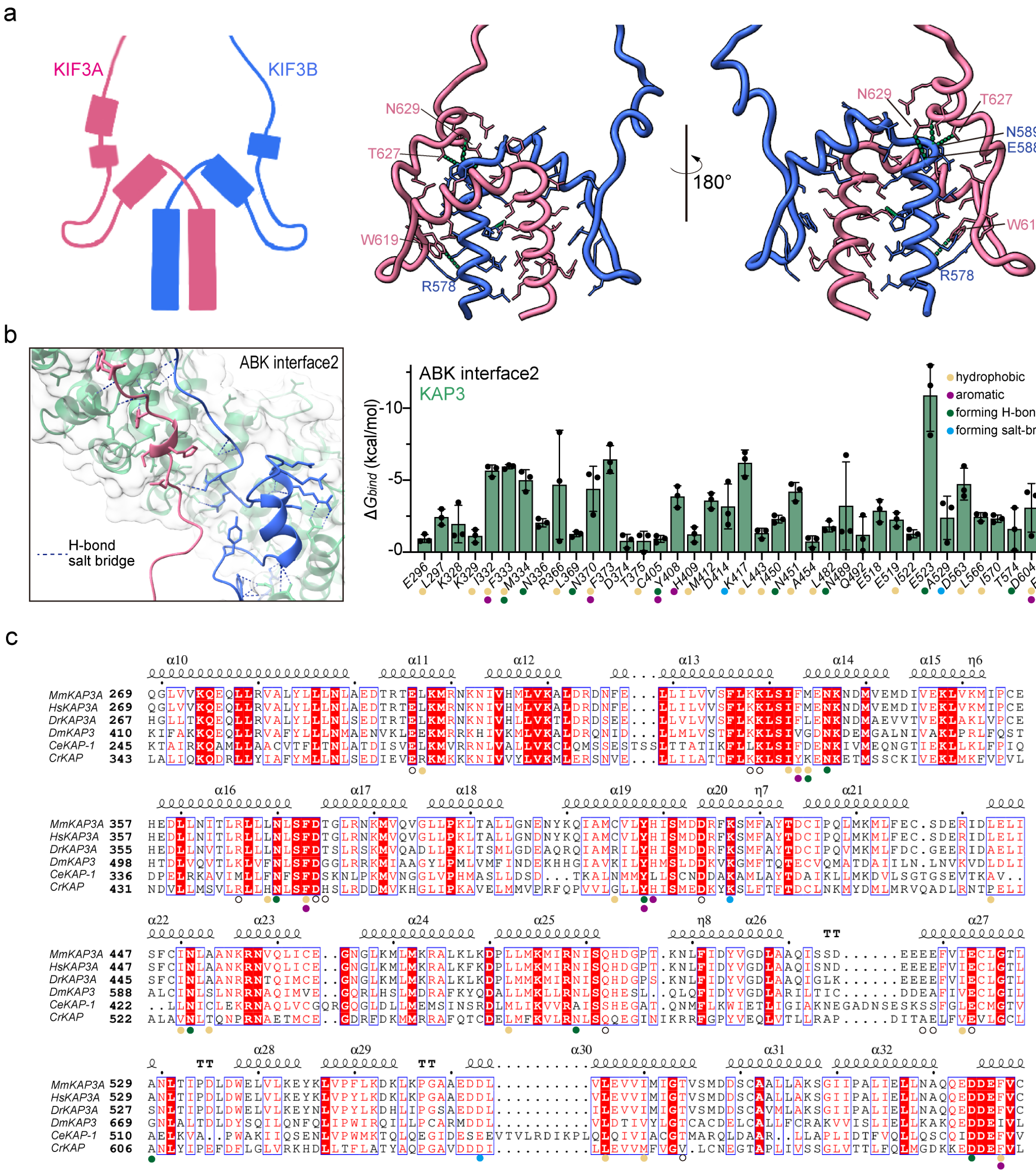


Supplementary Figure 3. KIF3A/B interaction and the binding contribution of KAP3 in ABK interface2.

**Fig. S3. KIF3A/B interaction and the binding contribution of KAP3 in ABK interface2.** (**a**) Ribbon representation of the KIF3A/B stalk-tail junction. Key interacting residues and hydrogen bonds are labeled. (**b**) Cartoon representation of the ABK interface2 structure and hydrogen bond network (left). Binding free energy contributions of KAP3 residues involved in the interface are summarized (right). Key interacting residues are labeled. (**c**) Multiple sequence alignment of KAP3 residues involved in ABK interface2 across species. Key interacting residues are labeled.

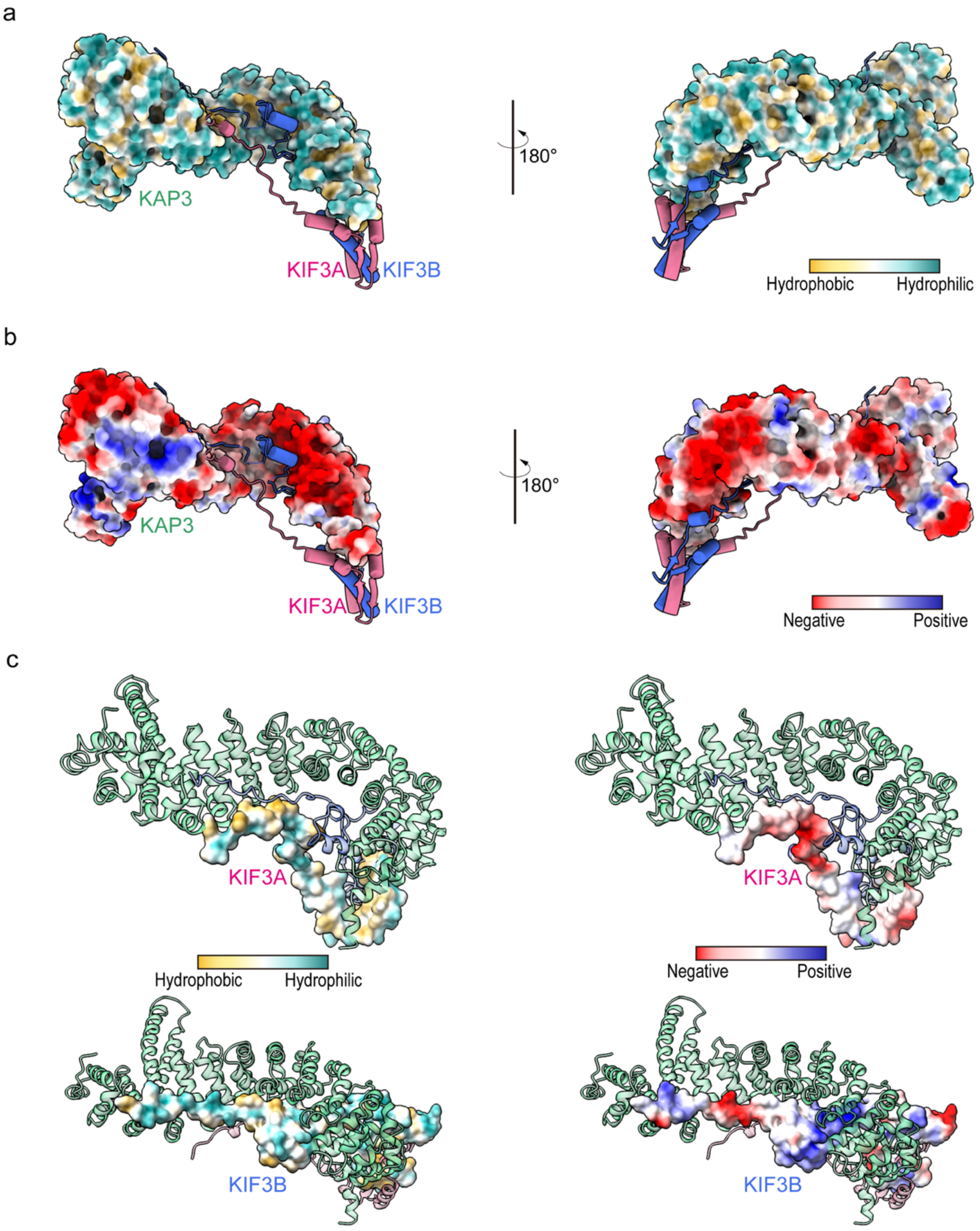


Supplementary Figure 4. Surface hydrophobicity and charge of KIF3A/B and KAP3

**Fig. S4. Surface hydrophobicity and charge of KIF3A/B and KAP3.** (**a**, **b**) Hydrophobic and electrostatic properties of the KAP3 surface, with KIF3 structures shown as cartoons. (**c**) Hydrophobic and electrostatic properties of the KIF3 surface, with KAP3 structures shown as cartoons.

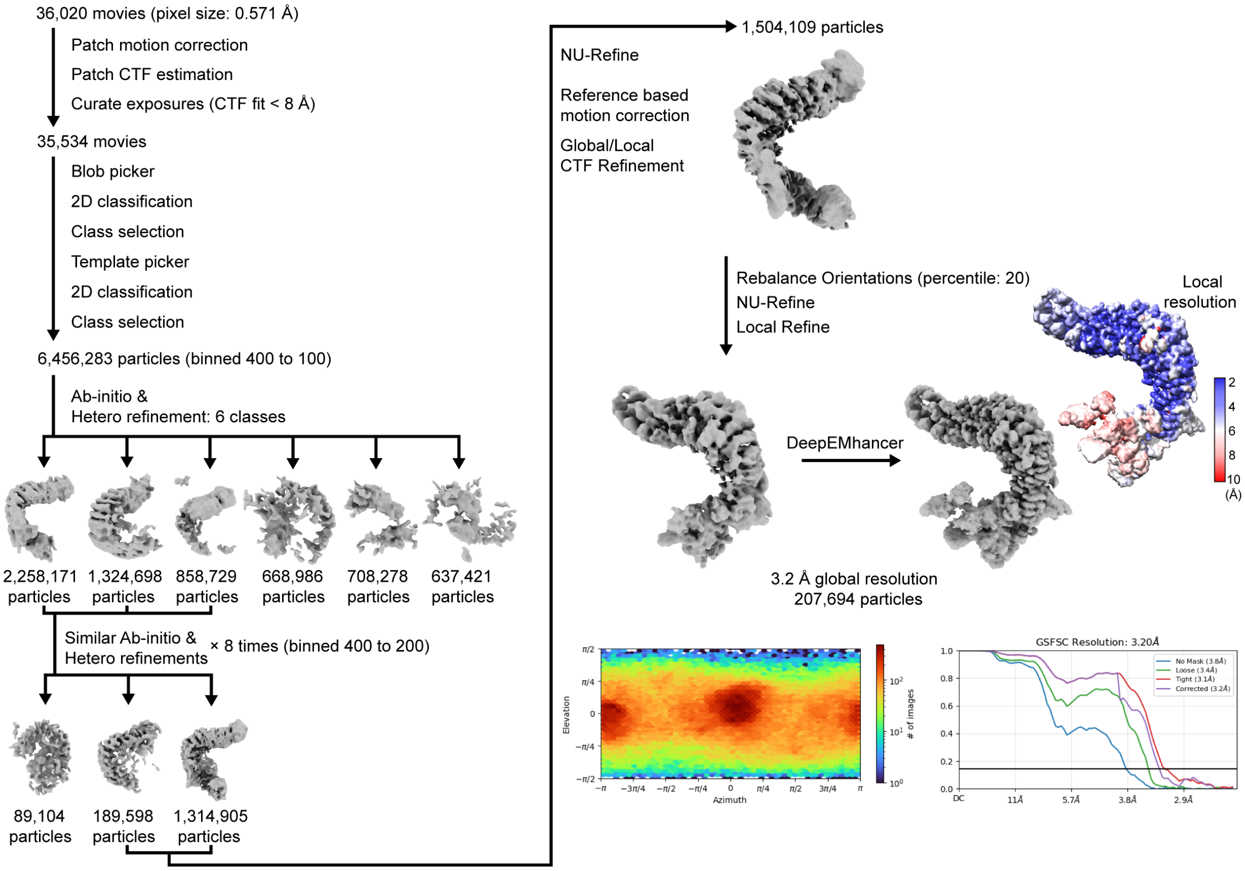


Supplementary Figure 5. Cryo-EM data processing for the KIF3A/B/KAP3-APCARM complex

**Fig. S5. Cryo-EM data processing workflow for the KIF3A/B/KAP3-APC complex.**

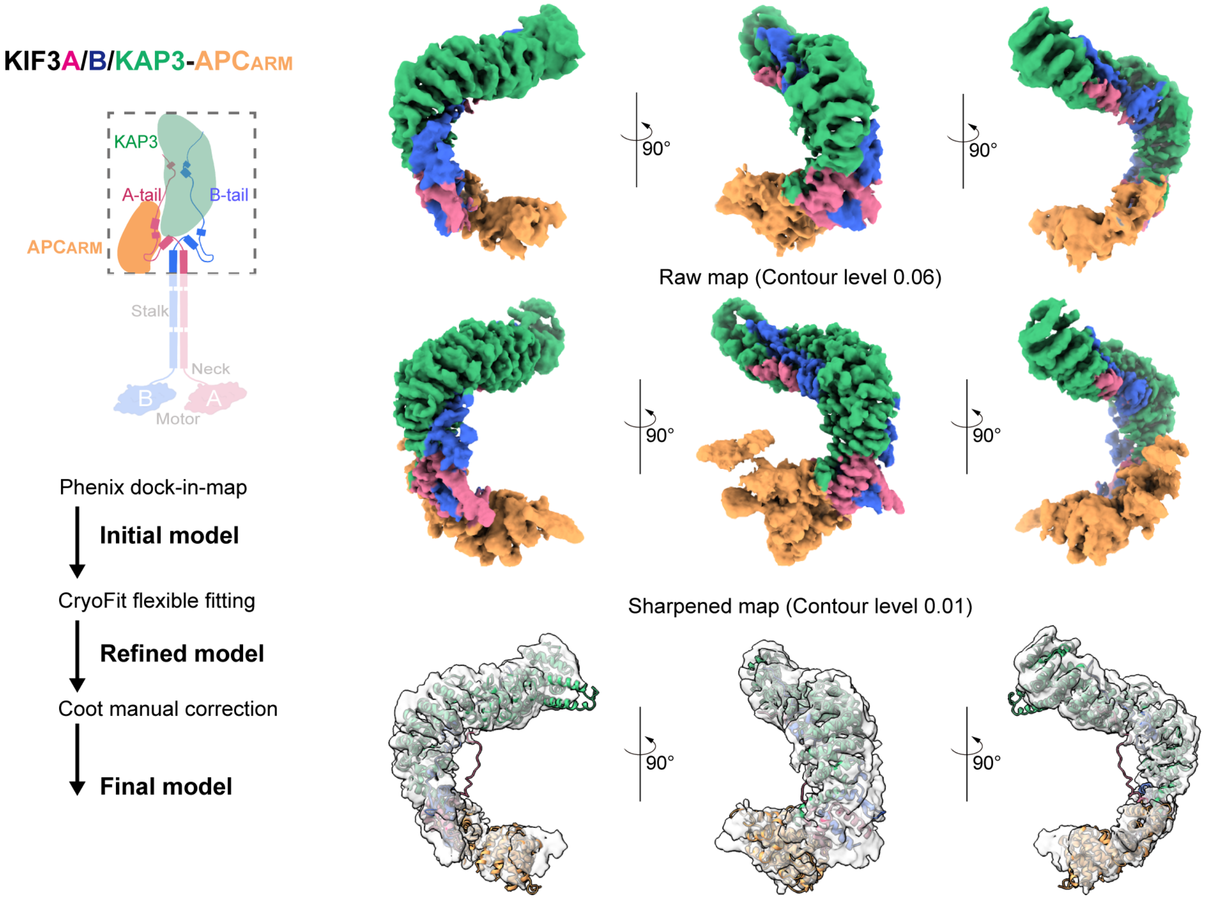


Supplementary Figure 6. Model building for the KIF3A/B/KAP3-APCARM complex

**Fig. S6. Model building for the KIF3A/B/KAP3-APC complex.**

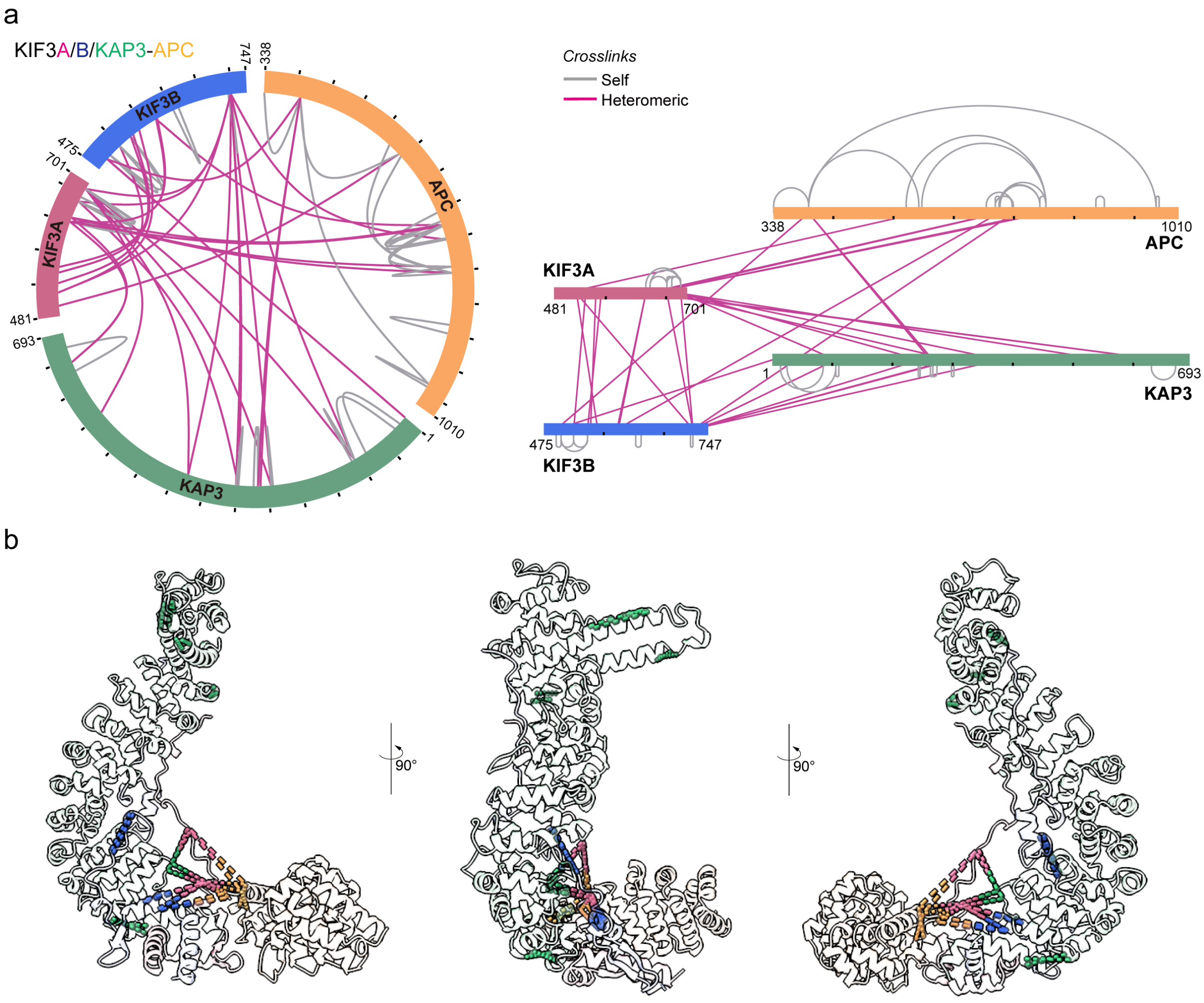


Supplementary Figure 7. Cross-linking mass spectrometry (XL-MS) results of the KIF3A/B/KAP3-APC complex

**Fig. S7. Crosslinking mass spectrometry (XL-MS) results of the KIF3A/B/KAP3-APC complex.** (**a**) Circular representation (left) and 2D diagram (right) of XL-MS results for the KIF3 C-terminal/KAP3-APC complex on the constructs. Intermolecular crosslinks are shown in purple, and intramolecular crosslinks are shown in gray. Only crosslinks visible in the structural model are displayed. Full data are available in fig. S7 and table S2. (**b**) XL-MS results mapped onto the structural model. Intermolecular crosslinks are depicted using dashed lines in two colors, each corresponding to the respective components, while intramolecular crosslinks are represented by dashed lines in a single color matching the component. Crosslink pairs are within theoretical distances.

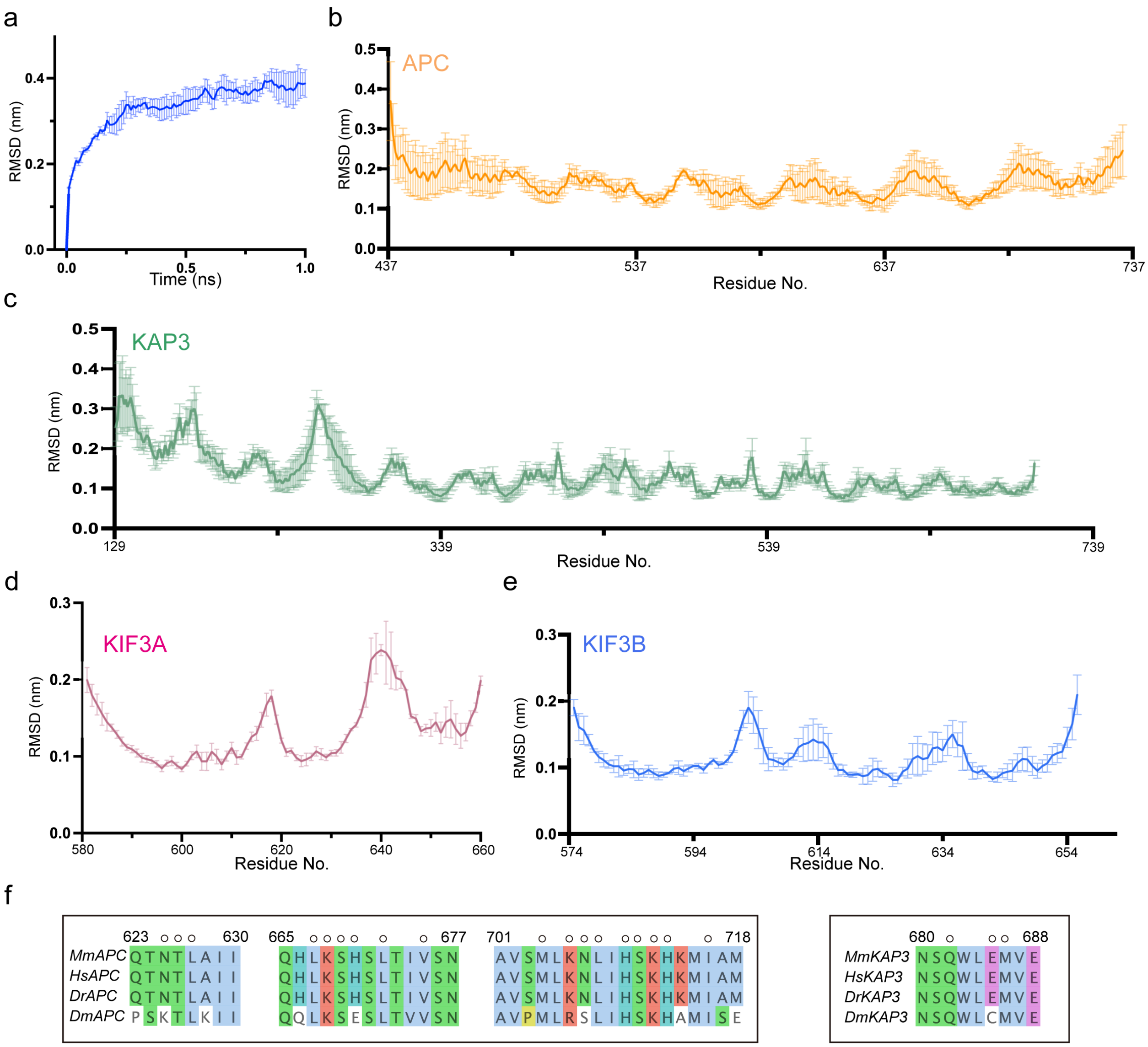


Supplementary Figure 8. Molecular dynamics and MMPBSA analysis of the KIF3A/B/KAP3-APC complex

**Fig. S8. Molecular dynamics and MMPBSA analysis of the KIF3A/B/KAP3-APC complex.**

(**a**) RMSD changes during three replicate MD simulations. (**b–e**) RMSD changes of APC, KAP3, KIF3A, and KIF3B residues before and after MD simulations. (**f**) Multiple sequence alignment of KAP3 and APC residues involved in KIF3/KAP3-APC binding across species. Key interacting residues are labeled. The alignment was generated using Clustal Omega (https://www.ebi.ac.uk/jdispatcher/msa/clustalo).

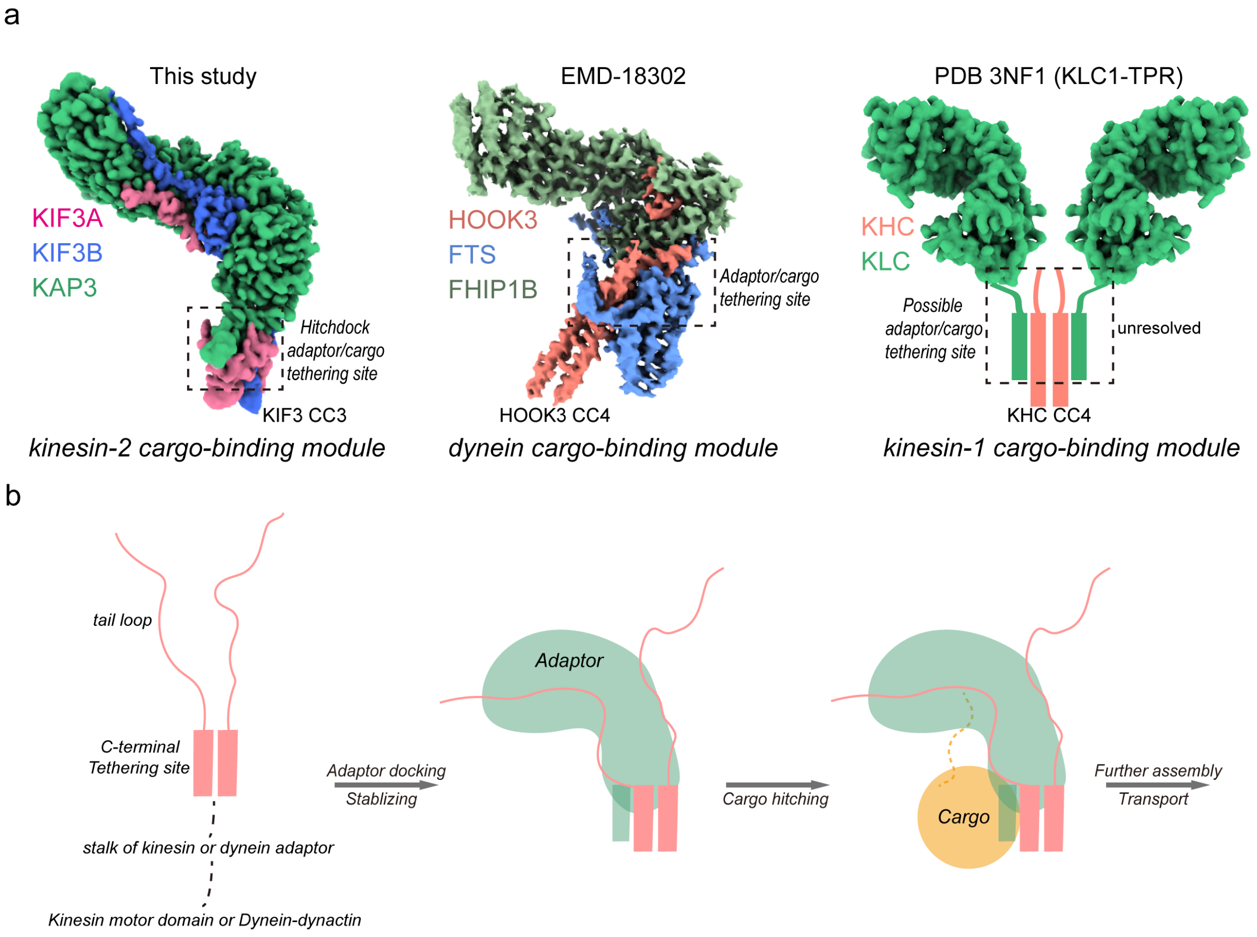


Supplementary Figure 9. Hook-like structures facilitate cargo recognition for kinesins and dyneins

**Fig. S9. Hook-like cargo-binding structures suggest a shared mechanism in kinesin and dynein cargo/adaptor assembly.** (**a**) Comparison of the KIF3/KAP3 cargo-binding module map obtained in this study with the dynein FTS-HOOK-FHIP1B (FTF) cargo-binding adaptor complex map and a possible kinesin-1 KHC/KLC model. The FTF density is derived from (EMDB: EMD-18302), and the density of the KLC1 TPR domain is generated based on its crystal structure (PDB: 3NF1). (**b**) A proposed common model for kinesin/dynein cargo recognition mechanism. The C-terminus of kinesin 1/2 and dynein adaptors contains an adaptor/cargo tethering site formed by helices, which mediates adaptor assembly and further recognition and binding to cargo. The form and contribution of the tethering site in cargo binding may differ; for kinesin-1 and dynein, cargo specificity is provided by the cargo-adaptor protein interaction (dashed line), while for kinesin-2, it is provided by the tethering site itself.

**Table S1. Cryo-EM data collection, modeling, and refinement statistics.**

| | KIF3A/B/KAP3 | KIF3A/B/KAP3-APCARM |
|---|---|---|
| **Data collection and processing** | | |
| Microscopy | CRYO ARM 200 | CRYO ARM 200 |
| Detector | Gatan K3 | Gatan K3 |
| Voltage | 200 kV | 200 kV |
| Cs (mm) | 1.55 | 1.55 |
| Slit width (eV) | 20 | 20 |
| Defocus range (μm) | 0.6 ; 1.8 | 0.6 ; 1.8 |
| Electron exposure (*e/Å2)* | 65.3 | 64.7 |
| Pixel size (*Å*) | 0.571 | 0.571 |
| Fractions (no.) | 75 | 80 |
| Movies collected/used | 34,176/33,576 | 36,020/35,534 |
| Single particles (no.) | 9,701,817 | 6,456,283 |
| Single particles used (no.) | 539,244 (class1) / 665,445 (class2) | 207,694 |
| Overall resolution (*Å*) | 2.83 (class1) / 2.75 (class2) | 3.20 |
| FSC threshold | 0.143 | 0.143 |
| **Model building and refinement** | | |
| Model composition | | |
| Non-hydrogen atoms | 6092 | 1035 |
| Protein residues | 742 | 1121 |
| R.m.s. deviation | | |
| Bond lengths (*Å*) | 0.006 | 0.008 |
| Bond angles (°) | 0.837 | 1.370 |
| Validation | | |
| MolProbity score | 2.38 | 2.35 |
| Clashscore | 15.68 | 22.29 |
| Poor rotamers (%) | 2.64 | 1.79 |
| Ramachandran plot | | |
| Favored (%) | 94.70 | 95.50 |
| Allowed (%) | 5.30 | 4.05 |
| Outliers (%) | 0 | 0.45 |

### Table S2. Cross-links detected by cross-linking mass spectrometry.

| Modified sequence1 | Modified sequence2 | Score | Crosslink type | Protein1 | Protein2 | Linked residue pairs |
|---|---|---|---|---|---|---|
| _AEEQEKLLEESNMELEER_ | _KESTLK_ | 142.1 | Inter-protein | KIF3A(P28741) | APC(Q61315) | K497-K579 |
| _VWTMLMAAKSEMADLQQEHQR_ | _LFSKLQAVK_ | 199.49 | Inter-protein | KIF3A(P28741) | KIF3B(Q61711) | K559-K547 |
| _VWTM(Oxidation(M))LMAAKSEMADLQQEHQR_ | _LFSKLQAVK_ | 64.723 | Inter-protein | KIF3A(P28741) | KIF3B(Q61711) | K559-K547 |
| _KQTPVPDK_ | _NKIMNR_ | 182.13 | Inter-protein | KIF3A(P28741) | KIF3B(Q61711) | K633-K598 |
| _QSLMKLER_ | _TELKMR_ | 62.2 | Inter-protein | KIF3A(P28741) | KAP3(P70188) | K668-K298 |
| _QSLMKLER_ | _PRTSKGK_ | 30.612 | Intra-protein | KIF3A(P28741) | KIF3A(P28741) | K668-K676 |
| _QSLMKLERPR_ | _TELKMR_ | 64.654 | Inter-protein | KIF3A(P28741) | KAP3(P70188) | K668-K298 |
| _QSLMKLERPR_ | _QTPVPDKK_ | 47.288 | Intra-protein | KIF3A(P28741) | KIF3A(P28741) | K668-K640 |
| _RSAKPETVIDSLLQ_ | _TSKGKAR_ | 77.662 | Intra-protein | KIF3A(P28741) | KIF3A(P28741) | K691-K676 |
| _SAKPETVIDSLLQ_ | _NKMVQVGLLPK_ | 108.71 | Inter-protein | KIF3A(P28741) | KAP3(P70188) | K691-K380 |
| _SAKPETVIDSLLQ_ | _KAVDEDLENQTLR_ | 95.015 | Inter-protein | KIF3A(P28741) | KAP3(P70188) | K691-K246 |
| _SAKPETVIDSLLQ_ | _KQTPVPDK_ | 89.959 | Intra-protein | KIF3A(P28741) | KIF3A(P28741) | K691-K633 |
| _SAKPETVIDSLLQ_ | _DSLPGKEK_ | 70.332 | Inter-protein | KIF3A(P28741) | KAP3(P70188) | K691-K106 |
| _SAKPETVIDSLLQ_ | _QSLMKLER_ | 66.004 | Intra-protein | KIF3A(P28741) | KIF3A(P28741) | K691-K668 |
| _SAKPETVIDSLLQ_ | _LVPFLKDK_ | 46.746 | Inter-protein | KIF3A(P28741) | KAP3(P70188) | K691-K552 |
| _SAKPETVIDSLLQ_ | _QTPVPDKK_ | 35.891 | Intra-protein | KIF3A(P28741) | KIF3A(P28741) | K691-K640 |
| _MQGEDAR_ | _ILEQKR_ | 30.57 | Inter-protein | KAP3(P70188) | KIF3B(Q61711) | M1-K495 |
| _GGNIDVHPSEKALIVQYEVEATILGEMGDPMLGER | _DSLPGKEK_ | 75.141 | Intra-protein | KAP3(P70188) | KAP3(P70188) | K25-K106 |
| _LIHPSKLSEVEQLLYYLQNR | _VKGGNIDVHPSEK | 122.53 | Intra-protein | KAP3(P70188) | KAP3(P70188) | K85-K14 |
| _LIHPSKLSEVEQLLYYLQNR_ | _RKVKGGNIDVHPSEK_ | 63.573 | Intra-protein | KAP3(P70188) | KAP3(P70188) | K85-K14 |
| _RDSLPGKEK_ | _KEKSSK_ | 21.993 | Intra-protein | KAP3(P70188) | KAP3(P70188) | K106-K111 |
| _KAVDEDLENQTLR_ | _HELWQEELSKK_ | 229.68 | Intra-protein | KAP3(P70188) | KAP3(P70188) | K246-K243 |
| _KAVDEDLENQTLR_ | _RHELWQEELSKK_ | 97.836 | Intra-protein | KAP3(P70188) | KAP3(P70188) | K246-K243 |
| _KAVDEDLENQTLR_ | _SAKPETVIDSLLQ_ | 71.085 | Inter-protein | KAP3(P70188) | KIF3A(P28741) | K246-K691 |
| _KYQGLVVK_ | _DYDKTFK_ | 183.24 | Intra-protein | KAP3(P70188) | KAP3(P70188) | K267-K263 |
| _YQGLVVKQEQLLR_ | _DYDKTFK_ | 60.628 | Intra-protein | KAP3(P70188) | KAP3(P70188) | K274-K263 |
| _NKNIVHMLVK_ | _TELKMR_ | 220.51 | Intra-protein | KAP3(P70188) | KAP3(P70188) | K302-K298 |
| _NKNIVHMLVK_ | _TELKM(Oxidation(M))R_ | 169.12 | Intra-protein | KAP3(P70188) | KAP3(P70188) | K302-K298 |
| _NKNIVHM(Oxidation (M))LVK_ | _TELKMR_ | 133.57 | Intra-protein | KAP3(P70188) | KAP3(P70188) | K302-K298 |
| _DVIIKETQAPAYLIDLMHDK_ | _KIQSEK_ | 191.49 | Intra-protein | KAP3(P70188) | KAP3(P70188) | K630-K670 |
| _ETQAPAYLIDLMHDKNNEIR_ | _KQTPVPDK_ | 77.505 | Inter-protein | KAP3(P70188) | KIF3A(P28741) | K645-K633 |
| _ETQAPAYLIDLMHDKNNEIR_ | _QTPVPDKK_ | 56.238 | Inter-protein | KAP3(P70188) | KIF3A(P28741) | K645-K640 |
| _ASAALHNIIHSQPDDKR_ | _DYDKTFK_ | 141.46 | Inter-protein | APC(Q61315) | KAP3(P70188) | K396-K263 |
| _ASAALHNIIHSQPDDKR_ | _KESTLK_ | 141.32 | Intra-protein | APC(Q61315) | APC(Q61315) | K396-K579 |
| _ASAALHNIIHSQPDDKR_ | _ILEQKR_ | 116.51 | Inter-protein | APC(Q61315) | KIF3B(Q61711) | K396-K495 |
| _ASAALHNIIHSQPDDKR_ | _KYQGLVVK_ | 79.346 | Inter-protein | APC(Q61315) | KAP3(P70188) | K396-K338 |
| _ASAALHNIIHSQPDDKR_ | _ASAALHNIIHSQPDDKR_ | 37.013 | Intra-protein | APC(Q61315) | APC(Q61315) | K396-K267 |

| | | | | | | |
|---|---|---|---|---|---|---|
| _ESTLKSVLSALWN LSAHCTENK_ | _HKQNLYGDYAF DANR_ | 17.619 | Intra-protein | APC(Q61315) | APC(Q61315) | K584-K790 |
| _NPKDQEALWDMG AVSMLK_ | _NLMANRPAKYK _ | 129.05 | Intra-protein | APC(Q61315) | APC(Q61315) | K691-K734 |
| _DQEALWDMGAVS MLKNLIHSK_ | _YKDANIMSPGSS LPSLHVR_ | 71.085 | Intra-protein | APC(Q61315) | APC(Q61315) | K706-K736 |
| _DQEALWDMGAVS MLKNLIHSK_ | _HKMIAMGSAAA LR_ | 65.207 | Intra-protein | APC(Q61315) | APC(Q61315) | K706-K714 |
| _NLIHSKHK_ | _KQTPVPDK_ | 122.52 | Inter-protein | APC(Q61315) | KIF3A(P28741) | K712-K633 |
| _HKMIAMGSAAAL R_ | _KQTPVPDK_ | 134.49 | Inter-protein | APC(Q61315) | KIF3A(P28741) | K714-K633 |
| _HKMIAMGSAAAL R_ | _QTPVPDKK_ | 101.6 | Inter-protein | APC(Q61315) | KIF3A(P28741) | K714-K640 |
| _ALEAELDAQHLSE TFDNIDNLSPKASH R_ | _HKMIAMGSAAA LR_ | 209.38 | Intra-protein | APC(Q61315) | APC(Q61315) | K780-K714 |
| _ALEAELDAQHLSE TFDNIDNLSPKASH R_ | _HKQNLYGDYAF DANR_ | 158.94 | Intra-protein | APC(Q61315) | APC(Q61315) | K780-K790 |
| _ALEAELDAQHLSE TFDNIDNLSPKASH R_ | _QTPVPDKK_ | 95.138 | Inter-protein | APC(Q61315) | KIF3A(P28741) | K780-K640 |
| _ALEAELDAQHLSE TFDNIDNLSPKASH R_ | _HKM(Oxidation (M))IAMGSAAAL R_ | 77.51 | Intra-protein | APC(Q61315) | APC(Q61315) | K780-K714 |
| _ALEAELDAQHLSE TFDNIDNLSPKASH R_ | _HKMIAM(Oxidati on (M))GSAAALR_ | 50.492 | Intra-protein | APC(Q61315) | APC(Q61315) | K780-K714 |
| _ALEAELDAQHLSE TFDNIDNLSPKASH R_ | _NLIHSKHK_ | 37.44 | Intra-protein | APC(Q61315) | APC(Q61315) | K780-K712 |
| _HKQNLYGDYAFD ANR_ | _HKMIAMGSAAA LR_ | 160.47 | Intra-protein | APC(Q61315) | APC(Q61315) | K790-K714 |
| _HKQNLYGDYAFD ANR_ | _QTPVPDKK_ | 90.334 | Inter-protein | APC(Q61315) | KIF3A(P28741) | K790-K640 |
| _HKQNLYGDYAFD ANR_ | _ADVNSKK_ | 67.115 | Intra-protein | APC(Q61315) | APC(Q61315) | K790-K558 |
| _HKQNLYGDYAFD ANR_ | _HKM(Oxidation(M ))IAMGSAAALR_ | 48.288 | Intra-protein | APC(Q61315) | APC(Q61315) | K790-K714 |
| _GLQITTTAAQIAK VMEEVSAIHTSQDD R_ | _GIGLSAYHPTTE NAGTSSKR_ | 81.179 | Intra-protein | APC(Q61315) | APC(Q61315) | K887-K873 |
| _SSNDSLNSVTSSD GYGKR_ | _GQMKPSVESYS EDDESK_ | 85.884 | Intra-protein | APC(Q61315) | APC(Q61315) | K973-K978 |
| _SSNDSLNSVTSSD GYGKR_ | _ASAALHNIIHSQ PDDKR_ | 34.474 | Intra-protein | APC(Q61315) | APC(Q61315) | K973-K396 |
| _NIVDHTNEQQKIL EQKR_ | _QEIAEQKRR_ | 64.565 | Intra-protein | KIF3B(Q61711) | KIF3B(Q61711) | K495-K503 |
| QEIAEQKR | ILEQKR | 175.75 | Intra-protein | KIF3B(Q61711) | KIF3B(Q61711) | K503-K495 |
| _DEETLELKETYTS LQQEVDIK_ | _LDIEEKYTSLQE EAQGK_ | 145.13 | Inter-protein | KIF3B(Q61711) | KIF3A(P28741) | K524-K533 |
| _DEETLELKETYTS LQQEVDIK | _QEIAEQKR_ | 103.66 | Intra-protein | KIF3B(Q61711) | KIF3B(Q61711) | K524-K503 |
| _DEETLELKETYTS LQQEVDIK_ | _LFSKLQAVK_ | 98.165 | Intra-protein | KIF3B(Q61711) | KIF3B(Q61711) | K524-K547 |
| _LFSKLQAVK_ | _QEIAEQKR_ | 55.353 | Intra-protein | KIF3B(Q61711) | KIF3B(Q61711) | K547-K503 |
| _LQAVKAEIHDLQE EHIK_ | _KVWTMLMAAK _ | 188.31 | Inter-protein | KIF3B(Q61711) | KIF3A(P28741) | K552-K550 |
| _AEIHDLQEEHIKER _ | _KELEEK_ | 125.97 | Inter-protein | KIF3B(Q61711) | KIF3A(P28741) | K564-K518 |
| _HLIIENFIPLEEKNK _ | _KQTPVPDK_ | 110.92 | Inter-protein | KIF3B(Q61711) | KIF3A(P28741) | K596-K633 |
| _HLIIENFIPLEEKNK _ | _NLIHSKHK_ | 96.591 | Inter-protein | KIF3B(Q61711) | APC(Q61315) | K596-K712 |
| PVSAVGYKR | LENQQMMKR | 37.18 | Intra-protein | KIF3B(Q61711) | KIF3B(Q61711) | K636-K627 |
| _KSGSSSSSSGNPAS QFYPQSR_ | _KAVDEDLENQT LR_ | 145.75 | Inter-protein | KIF3B(Q61711) | KAP3(P70188) | K722-K246 |

| | | | | | | |
|---|---|---|---|---|---|---|
| _KSGSSSSSSGNPAS QFYPQSR_ | _KELEEK_ | 122.75 | Inter-protein | KIF3B(Q61711) | KIF3A(P28741) | K722-K518 |
| _KSGSSSSSSGNPAS QFYPQSR_ | _NKMVQVGLLPK _ | 107.45 | Inter-protein | KIF3B(Q61711) | KAP3(P70188) | K722-K380 |
| _KSGSSSSSSGNPAS QFYPQSR_ | _TELKMR_ | 106.28 | Inter-protein | KIF3B(Q61711) | KAP3(P70188) | K722-K298 |
| _KSGSSSSSSGNPAS QFYPQSR_ | _ARPKSGR_ | 94.203 | Intra-protein | KIF3B(Q61711) | KIF3B(Q61711) | K722-K718 |
| _KSGSSSSSSGNPAS QFYPQSR_ | _QSLMKLERPR_ | 75.533 | Inter-protein | KIF3B(Q61711) | KIF3A(P28741) | K722-K668 |
| _KSGSSSSSSGNPAS QFYPQSR_ | _DSLPGKEK_ | 72.421 | Inter-protein | KIF3B(Q61711) | KAP3(P70188) | K722-K106 |
| _KSGSSSSSSGNPAS QFYPQSR_ | _NKNIVHMLVK_ | 67.251 | Inter-protein | KIF3B(Q61711) | KAP3(P70188) | K722-K302 |
| _KSGSSSSSSGNPAS QFYPQSR_ | _SAKPETVIDSLL Q_ | 58.373 | Inter-protein | KIF3B(Q61711) | KIF3A(P28741) | K722-K691 |
| _KSGSSSSSSGNPAS QFYPQSR_ | _YKDANIMSPGSS LPSLHVR_ | 49.101 | Inter-protein | KIF3B(Q61711) | APC(Q61315) | K722-K736 |
